# ZBED2 is an antagonist of Interferon Regulatory Factor 1 and modulates cell identity in pancreatic cancer

**DOI:** 10.1101/868141

**Authors:** Tim D.D. Somerville, Yali Xu, Xiaoli S. Wu, Christopher R. Vakoc

## Abstract

Lineage plasticity is a prominent feature of pancreatic ductal adenocarcinoma (PDA) cells, which can occur via deregulation of lineage-specifying transcription factors. Here, we show that the zinc finger protein ZBED2 is aberrantly expressed in PDA and alters tumor cell identity in this disease. Unexpectedly, our epigenomic experiments reveal that ZBED2 is a sequence-specific transcriptional repressor of interferon-stimulated genes, which occurs through antagonism of Interferon Regulatory Factor 1 (IRF1)-mediated transcriptional activation at co-occupied promoter elements. Consequently, ZBED2 attenuates the transcriptional output and growth arrest phenotypes downstream of interferon signaling in multiple PDA cell line models. We also found that ZBED2 is preferentially expressed in the squamous molecular subtype of human PDA, in association with inferior patient survival outcomes. Consistent with this observation, we show that ZBED2 can repress the pancreatic progenitor transcriptional program, enhance motility, and promote invasion in PDA cells. Collectively, our findings suggest that high ZBED2 expression is acquired during PDA progression to suppress the interferon response pathway and to promote lineage plasticity in this disease.

**SIGNIFICANCE STATEMENT:** Pancreatic ductal adenocarcinoma (PDA) is one of the most lethal human malignancies, attributed in part to lineage infidelity downstream of deregulated lineage-specifying transcription factors (TFs). Here we define the biological effects of a poorly understood TF ZBED2 in the most aggressive subtype of PDA, defined by the expression of squamous lineage markers. Our study reveals two molecular functions of ZBED2 in PDA cells: an inhibitor of interferon response genes and a modifier of epithelial lineage programs. Both functions can be explained by the ability of ZBED2 to antagonize the functional output of Interferon Regulatory Factor-1 (IRF1). Our study reinforces the concept of aberrant lineage identity in cancer and highlights an unexpected connection between interferon response pathways and squamous-subtype PDA.

## INTRODUCTION

Cells are capable of adopting different identities along a phenotypic spectrum, a process referred to as lineage plasticity. While lineage plasticity is a critical feature of normal development and during wound-healing responses, it is also a powerful contributor to the pathogenesis of human cancer (1, 2). Through a multitude of genetic and non-genetic mechanisms, cancer cells gain access to diverse lineage and developmental transcriptional programs, resulting in heterogeneous cell identities emerging in a clonally-derived tumor. The consequences of lineage plasticity in human cancer are far-reaching, but include a prominent role in the acquisition of metastatic traits and in the evasion of targeted therapy (1).

Pancreatic ductal adenocarcinoma (PDA) is an emerging paradigm for the study of lineage plasticity in human cancer. While PDA is defined by its histopathological resemblance to ductal epithelial cells of the exocrine pancreas, studies in mice have shown that acinar cells can serve as a cell-of-origin of this disease, which trans-differentiate into the ductal fate following acquisition of a *Kras* activating mutation (3–5). At later stages of tumor development, aberrant up-regulation or silencing of master regulator transcription factors (TFs) in PDA can lead to reprogramming of ductal identity towards that of other cell lineages, including mesenchymal (6–8), foregut endodermal (9), or squamous epithelial fates (10–12). While each of these lineage transitions are capable of promoting disease progression in experimental systems, only the presence of squamous characteristics correlates with a shorter overall survival in human PDA patients (13, 14). For this reason, the identification of mechanisms that promote squamous trans-differentiation in PDA has become an active area of investigation in recent years (10–13, 15, 16).

The interferon (IFN) transcriptional response is a conserved pathway that protects organisms from infectious pathogens and malignancy (17, 18). IFN pathway activation occurs via autocrine or paracrine IFN signalling that can be triggered in response to the detection of foreign nucleic acids as well as ectopically located self-DNA (18, 19). Whereas almost all cell types can produce type I IFNs (e.g. IFN-α and IFN-β), type II IFN (i.e. IFN-γ) production is restricted to a subset of activated immune cells (20). IFN pathway activation promotes the transcriptional induction of hundreds of IFN-stimulated genes (ISGs), which encode diverse proteins with anti-viral, anti-proliferative, and immunostimulatory functions (21). The key TFs that promote ISG induction belong to the Signal Transducer and Activator of Transcription (STAT) and IFN Regulatory Factor (IRF) families, which can bind in an IFN-inducible manner at the promoters of ISGs (22, 23). In the classical pathway, phosphorylation of STATs downstream of IFN receptor activation triggers a rapid ISG response (23). This primary response includes the STAT-dependent transcriptional activation of several genes encoding IRFs, which subsequently drive an amplifier circuit resulting in sustained ISG induction (22). Within this complex transcriptional response, IRF1 is a critical positive regulator required for the full range of overlapping target gene activation following type I or type II IFN pathway activation (22). IRF1 is a broadly acting antiviral effector and exhibits tumor suppressor functions in multiple cellular contexts (24, 25). With respect to PDA, prior studies have shown that IRF1 can promote a differentiated epithelial cell state and inhibit cell proliferation (26, 27).

The ZBED gene family encodes nine zinc finger-containing TFs in humans, which originated from a domesticated DNA transposase gene from an hAT transposable element (28). While lacking in transposase activity, human ZBED TFs instead retain their zinc finger domain to perform sequence-specific DNA binding and function as transcriptional regulators in a cell-type specific manner (29–31). Within this family, ZBED2 is one of the least understood members, owing in part to its recent evolution and lack of a mouse ortholog (28). A prior genomewide-association study identified *ZBED2* as a candidate locus influencing risk of smoking-induced pancreatic cancer (32). More recently, *ZBED2* was found to be highly expressed in the basal layer of the epidermis, where it plays a role in regulating keratinocyte differentiation (33). Another study identified *ZBED2* as a marker of T cell exhaustion in human CD8 T cells, although the function of ZBED2 was not investigated in this context (34). We are unaware of any prior study characterizing a transcriptional function for ZBED2 or its role in cancer.

Here we identify ZBED2 as one of the most aberrantly up-regulated TFs in human PDA. This prompted our characterization of the transcriptional function of ZBED2, which we demonstrate to be a sequence-specific transcriptional repressor. We show that the repression targets of ZBED2 are highly enriched for genes within the IFN response pathway. By interacting with ISG promoters, ZBED2 blocks the transcriptional output and growth arrest phenotypes caused by IRF1 activation downstream of IFN stimulation. We also provide evidence that ZBED2 is preferentially expressed in squamous-subtype PDA tumors and promotes loss of pancreatic progenitor cell identity in this context. Collectively, our study suggests that aberrant ZBED2 expression in PDA cells blocks the IFN response and alters epithelial cell identity in this disease.

## RESULTS

### Aberrant *ZBED2* expression in pancreatic ductal adenocarcinoma correlates with inferior patient survival outcomes

In this study, we sought to identify novel TFs that deregulate cell identity in PDA. By evaluating previously published transcriptome data comparing normal human pancreas and PDA (35), we identified *ZBED2* as among the most aberrantly up-regulated TF genes in tumor and metastatic lesions (Fig. 1*A* and *B*). Notably, the fold-induction of *ZBED2* was comparable to other aberrantly expressed TFs in PDA, *FOXA1* and *TP63*, which have established roles in disease progression (Fig. 1*A* and Dataset S1) (9, 10). In two additional independent transcriptome studies of human PDA tumors (13, 36), we validated high levels of *ZBED2* expression in 16-20% of primary patient samples (Fig. 1*C* and *D*). We further corroborated the aberrant expression pattern of *ZBED2* in ∼20% of established organoid cultures derived from human PDA tumors, while *ZBED2* was expressed at low levels in organoids derived from normal pancreatic epithelial cells (Fig. 1*E* and Dataset S2) (37). Considering the limited number of prior studies of ZBED2, these observations prompted us to investigate the molecular function of ZBED2 in PDA.

**Fig. 1.**
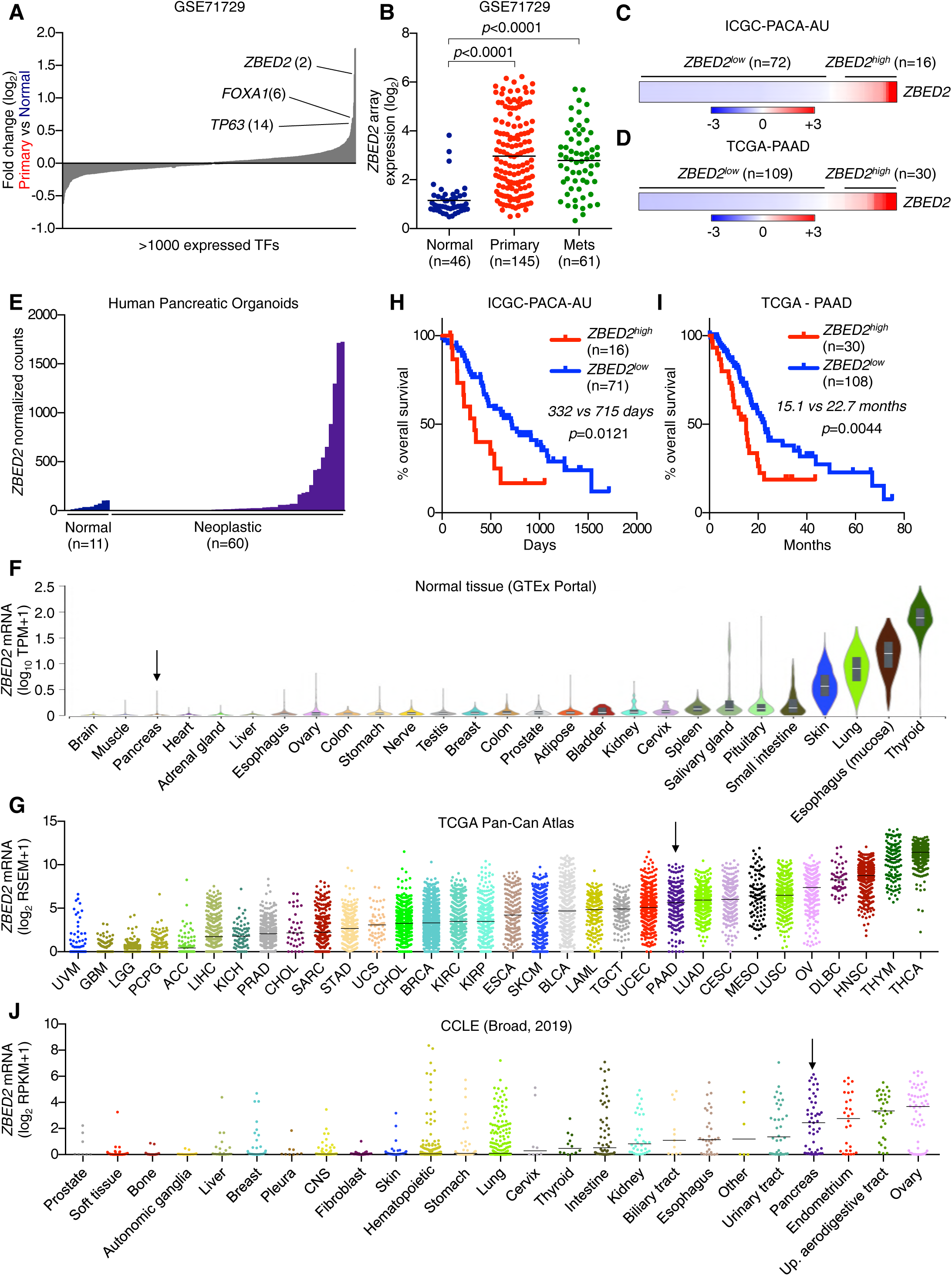
Aberrant *ZBED2* expression in pancreatic ductal adenocarcinoma correlates with inferior patient survival outcomes. (*A*) Expressed TFs ranked by their mean log_2_ fold change in primary PDA versus normal pancreas tissue samples. Selected transcription factors are labeled along with their rank. (*B*) *ZBED2* expression in normal pancreas tissue and primary and metastatic PDA tumor samples. *p* value was determined by one-way ANOVA. (*C* and *D*) *ZBED2* expression across PDA patient samples. Scale bar indicates the standardized expression value. (*E*) *ZBED2* expression in human organoids derived from normal pancreatic tissue or neoplastic PDA tumor samples. (*F* and *G*) *ZBED2* expression in normal tissue samples from the GTEx portal (*F*) or profiled tumors from the TCGA Pan-Cancer Atlas (*G*). Arrow indicates pancreas tissue. (*H* and *I*) Survival curves of patients stratified according to high or low *ZBED2* expression. *p* value was calculated using the log rank (Mantel-Cox) test. (*J*) *ZBED2* expression in cancer cell lines from the CCLE database. Arrow indicates pancreas tissue. See also Fig. S1.

To gain initial insight into ZBED2 function, we examined its expression across a diverse collection of normal and malignant human tissues using publicly available transcriptome data (38, 39). This analysis revealed a highly tissue-specific pattern of *ZBED2* expression in normal tissues, with skin, lung, esophageal mucosa, and thyroid tissues expressing *ZBED2* at the highest levels (Fig. 1*F*). In accord with the findings above, *ZBED2* expression was not detected in the normal human pancreas (Fig. 1*F*). Transcriptome analysis of 32 tumors types profiled by the TCGA Pan-Cancer Atlas (39) revealed pervasive *ZBED2* expression in numerous human cancer types (Fig. 1*G*). In specific malignancies, like thyroid carcinomas (THCA) and squamous cell carcinomas (HNSC, LUSC, ESCA), high *ZBED2* expression can be explained by its expression pattern in the normal tissue counterpart (i.e. cell-of-origin) of these tumors. However, in other cancer types, like PDA and ovarian cancer, *ZBED2* appears to be up-regulated in an aberrant manner (Fig. 1*F* and *G*). Remarkably, we found that patients with *ZBED2*^high^ tumors have a significantly shorter overall survival than patients with *ZBED2*^low^ tumors in the context of PDA, colorectal cancer, kidney cancer, glioblastoma/glioma, lung squamous cell carcinoma and lung adenocarcinoma (Fig. 1*H* and *I* and *SI Appendix*, Fig. S1*A*) (40). This highlights *ZBED2* as biomarker of disease aggressiveness across several tumor subtypes.

We further interrogated *ZBED2* expression in 1,156 cancer cell lines from the CCLE database (41) and identified high expression in ∼24% of cell lines (Fig. 1*J*). In accord with the TCGA analysis, *ZBED2*^high^ cell lines were significantly enriched for lines derived from pancreatic, ovarian, intestinal, endometrial and upper aerodigestive tract tumors (Fig. 1*J* and *SI Appendix*, Fig. S1*B*). In addition, *ZBED2*^high^ cell lines were significantly enriched for genetic alterations in *KRAS*, *TP53*, *CDKN2A* and *SMAD4* (*SI Appendix*, Fig. S1*C* and *D*), which are also the most common recurrently mutated genes in PDA (42–44). Despite this high level expression, analysis of CRISPR screening data from 559 cancer cell lines revealed that ZBED2 is not required to sustain cell growth in any cell line under standard tissue culture conditions (*SI Appendix*, Fig. S1*E*).

### ZBED2 inhibits the expression of genes in the interferon response pathway

To understand the cancer-relevant function of ZBED2, we performed RNA-seq analysis following *ZBED2* knockout or overexpression in PDA cell lines. Western blot analysis revealed high level expression of endogenous ZBED2 in three out of nine PDA cell lines examined (Fig. 2*A*). We used lentiviral CRISPR-Cas9 editing to target *ZBED2* in the highest expressing PDA cell line (PANC0403) with two independent sgRNAs and performed RNA-seq analysis. An unbiased gene set enrichment analysis (GSEA) of the *ZBED2* knockout RNA-seq data revealed induction of the IFN pathway as top-ranked gene signatures in this experiment (Fig. 2*B* and *C* and Dataset S3 and S4). To complement this approach, we performed RNA-seq after overexpressing a *ZBED2* cDNA in 15 different PDA cell lines, which likewise revealed consistent suppression of IFN pathway gene signatures as a top-ranked alteration (Fig. 2*B, D* and *E* and *SI Appendix*, Fig. S2*A* and Dataset S3-5). We confirmed the down-regulation of the ISGs *STAT2*, *CMPK2* and *MX1* following ectopic ZBED2 expression by RT-qPCR and western blotting (Fig. 2*F* and *SI Appendix*, Fig. S2*B*). We also confirmed the up-regulation of the ISGs *STAT2* and *CMPK2* in response to ZBED2 knockdown using shRNAs as an alternative approach to CRISPR (*SI Appendix*, Fig. S2*C* and *D*). Taken together, this transcriptome analysis implicates an inhibitory effect of ZBED2 on the IFN transcriptional response in PDA cells.

**Fig. 2.**
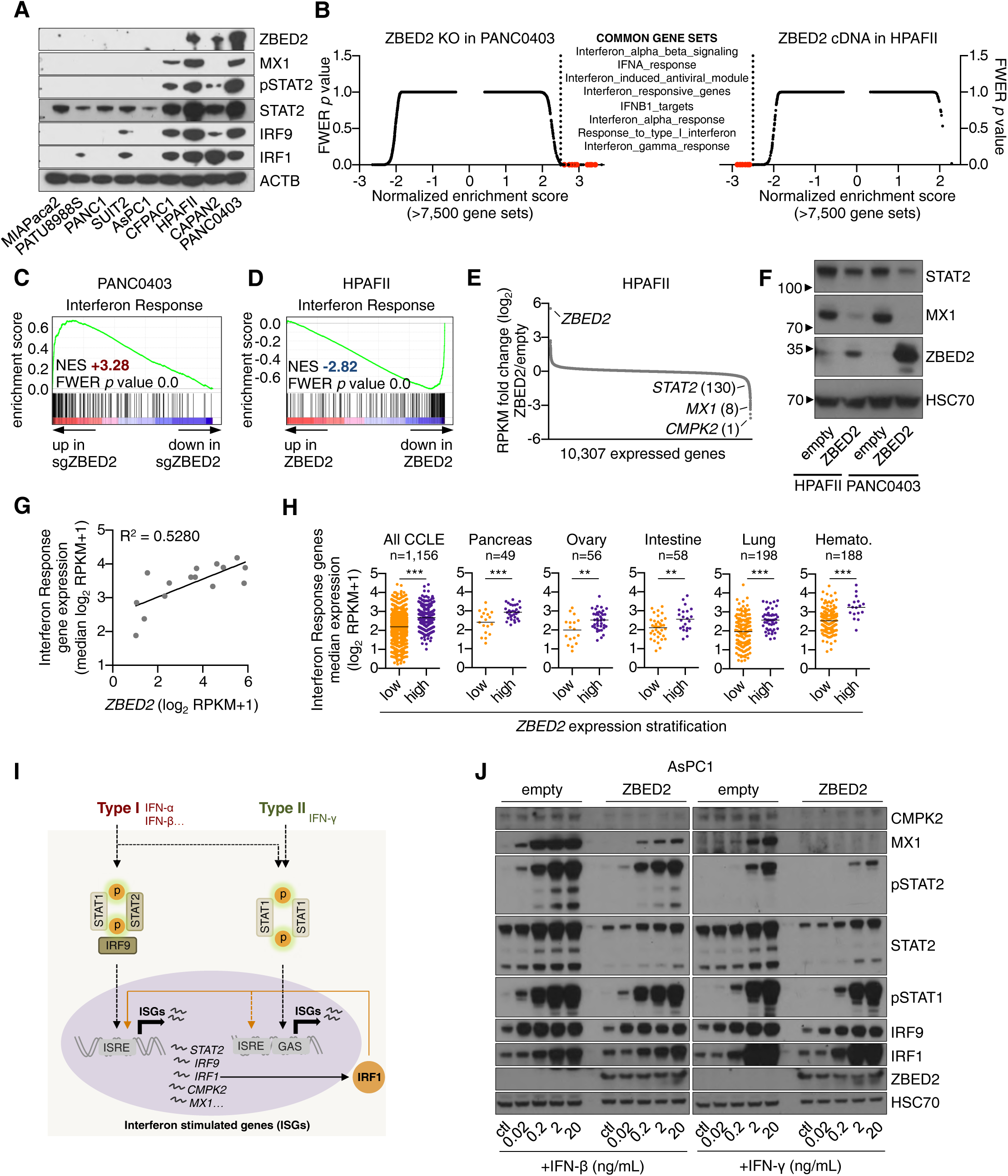
ZBED2 inhibits the expression of genes in the interferon response pathway. (*A*) Western blot analysis for ZBED2 and IFN pathway components in human PDA cell lines. (*B*) GSEA following ZBED2 knockout in PANC0403 cells (left panel) or ZBED2 cDNA expression in HPAFII cells (right panel) versus their respective controls. Normalized enrichment score (NES) and family-wise error rate (FWER) *p* value were ranked and plotted for 7,573 (PANC0403) and 7,593 (HPAFII) genes sets within MSigDB v7.0 (H1, C2, C5 and C6 collections). Each gene set is depicted as a single dot. (*C* and *D*) GSEA plots evaluating the Interferon Response signature upon ZBED2 knockout in PANC0403 cells (*C*) or ZBED2 cDNA expression in HPAFII cells (*D*). (*E*) Gene expression changes in HPAFII cells infected with ZBED2 cDNA versus those infected with an empty vector control. *ZBED2* and the ISGs *STAT2*, *MX1* and *CMPK2* are labeled along with their rank with respect to down regulated genes. (*F*) Western blot analysis for HSC70, ZBED2 and the IFN pathway proteins STAT2 and MX1 following ZBED2 cDNA expression in HPAFII and PANC0403 cells. (*G*) *ZBED2* expression versus the median expression value of Interferon Response genes across 15 human PDA cell lines. (*H*) Median expression values of Interferon Response genes across all cancer cell lines within the CCLE database (left panel) or in the indicated tissue lineages stratified according to high or low *ZBED2* expression. Each cell line is depicted as a single dot. **p <0.01, ***p <0.001 by Student’s t test. (*I*) Schematic representation of IFN signaling pathways and the IRF1 amplification loop. ISGs, interferon stimulated genes. Adapted from ref (22). (*J*) Western blot analysis for HSC70, ZBED2 and the indicated IFN pathway components following 12 hour stimulation with the increasing concentrations of IFN-β or IFN-γ in AsPC1 cells stably expressing ZBED2 or the empty vector. See also Fig. S2.

Despite the inhibitory effect of ZBED2 on IFN pathway genes, our western blot and RNA-seq analysis revealed that *ZBED2*^high^ PDA lines tended to have higher baseline level of IFN pathway activation, as judged by levels of STAT2 phosphorylation and ISG expression (Fig. 2*A* and *G*). This high basal activity of the IFN pathway was not attributed to autocrine IFN production, since inactivating the IFN alpha receptor (IFNAR1) did not suppress ISG levels (*SI Appendix*, Fig. S2*E* and *F*). Moreover, the positive correlation between levels of *ZBED2* and IFN pathway genes was observed across all 1,156 cell lines from the CCLE, encompassing PDA lines as well other diverse cancer types (Fig. 2*H* and *SI Appendix*, Fig. S2*G*). Additionally, we did not observe an upregulation of *ZBED2* following stimulation with IFN-β or IFN-γ, indicating that *ZBED2* is not itself an ISG in this context (*SI Appendix*, Fig. S2*H*). Taken together, these findings suggest that ZBED2 expression may have been acquired in PDA cells to dampen a pre-existing activation state of the IFN pathway.

We next validated the inhibitory effect of ZBED2 on the IFN pathway by performing western blot analysis of IFN components following exposure of PDA cells to IFN-β or IFN-γ (Fig. 2*I* and *J*). In these experiments, ZBED2 had no effect on STAT phosphorylation, which suggests that ZBED2 does not function in the signaling pathway downstream of receptor activation. In addition, this analysis revealed that specific ISG products were induced normally following IFN treatment (e.g. IRF1, IRF9), while others were attenuated by the presence of ZBED2 (STAT2, MX1, and CMPK2) (Fig. 2*J*). These findings suggest that ZBED2 functions as a repressor of a specific subset of ISGs.

### ChIP-seq analysis implicates ZBED2 as a sequence-specific repressor of ISG promoters

In order to investigate the function of ZBED2 at the molecular level, we performed chromatin immunoprecipitation followed by DNA sequencing (ChIP-seq) analysis of lentivirally expressed FLAG-ZBED2 in AsPC1 and SUIT2 PDA cell lines. We observed high concordance between ZBED2 occupancy in both cell lines, enabling us to define a set of 2,451 high confidence binding sites (Fig. 3*A* and *SI Appendix*, Fig. S3*A* and Dataset S6), which was highly biased towards promoter regions (Fig. 3*B*). Notably, genomic occupancy of ZBED2 was globally correlated with its repressive effect on transcription when comparing ChIP-seq with the aforementioned RNA-seq analyses. This included several ISGs (e.g. *CMPK2* and *STAT2*) that are both downregulated and located near ZBED2 binding sites (Fig. 3*C* and Dataset S7). To further verify the repressor function of ZBED2, we made use of GAL4 fusion protein to artificially tether ZBED2 to a plasmid with luciferase expression downstream of a thymidine kinase promoter. Unlike the established activator protein IRF1, tethering ZBED2 to this promoter resulted in a significant decrease in luciferase expression (Fig. 3*D*). Together, these findings suggest that ZBED2 is a transcriptional repressor that occupies specific sites in the genome.

**Fig. 3.**
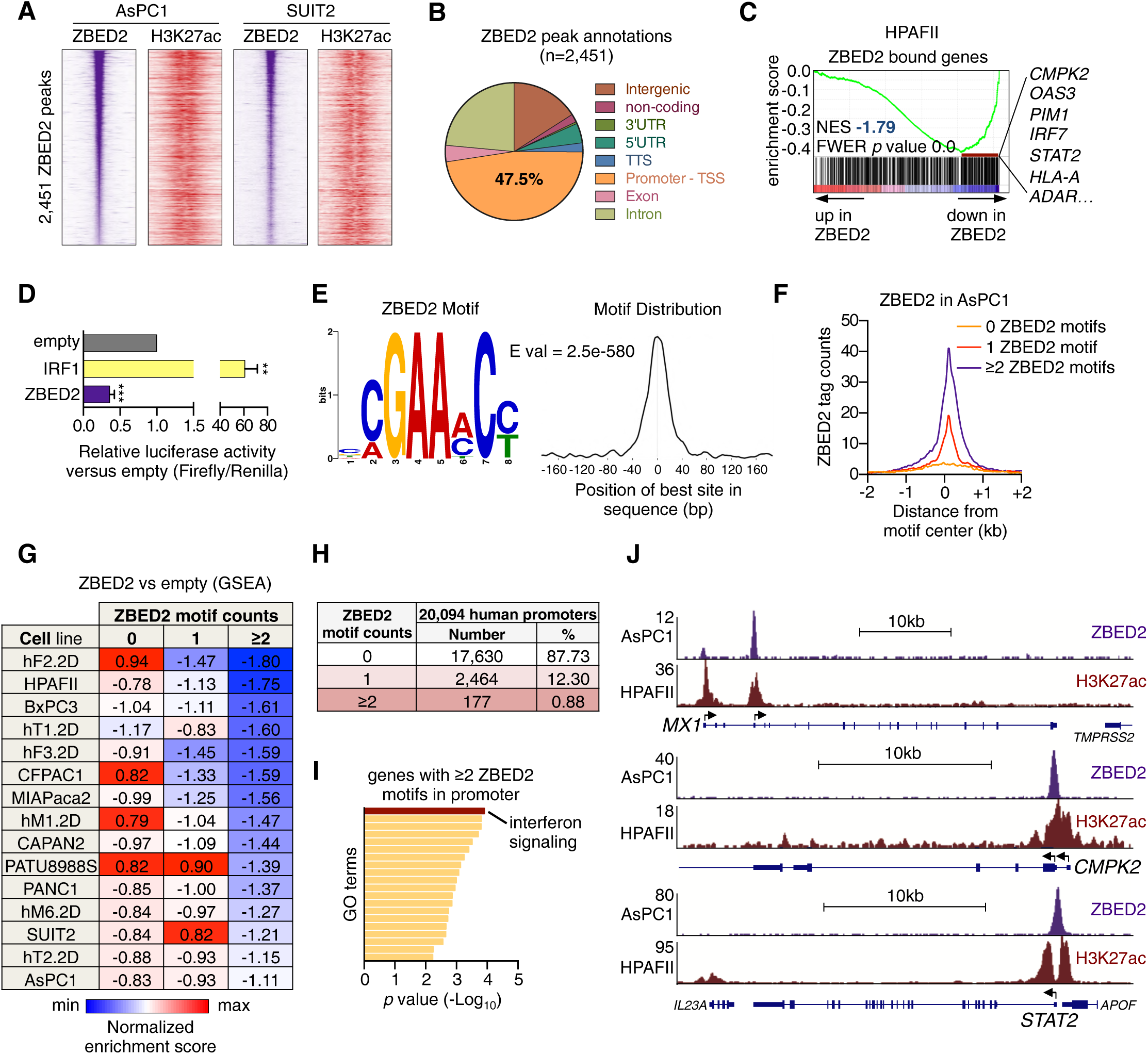
ChIP-seq analysis implicates ZBED2 as a sequence-specific repressor of ISG promoters. (*A*) Density plots showing FLAG-ZBED2 and H3K27ac enrichment surrounding a 2-kb interval centered on the summit of 2,451 high-confidence ZBED2 peaks in AsPC1 and SUIT2 cells, ranked by FLAG-ZBED2 peak intensity in AsPC1 cells. (*B*) Pie chart showing the distribution of high-confidence FLAG-ZBED2 peaks in AsPC1 cells. TTS, transcription termination site; TSS, transcription start site; UTR, untranslated region. (*C*) GSEA plot evaluating ZBED2 bound genes upon ZBED2 cDNA expression in HPAFII cells. Leading edge (indicated by the red line), Interferon Response genes are listed. (*D*) GAL4 fusion reporter assay testing full length ZBED2 and IRF1 transactivation activity normalized to Renilla luciferase internal control. Mean+SEM is shown. n=3. **p <0.01, ***p <0.001 by Student’s t test. (*E*) ZBED2 ChIP-seq derived *de novo* motif logo, distribution, and E value for the ZBED2 binding motif derived using MEME-ChIP. (*F*) Meta-profile comparing ZBED2 occupancy in AsPC1 cells around the center of peaks with 0, 1 or ≥2 motif counts. ZBED2 binding intensity is shown as sequencing depth normalized tag count. (*G*) Summary of GSEA evaluating ZBED2 bound genes at promoter regions with 0, 1 or ≥2 motif counts upon ZBED2 cDNA expression in 15 PDA cell lines. (*H*) Summary of ZBED2 motif density frequency in the human genome. The ZBED2 motif frequencies were calculated in all annotated human promoter regions (n=73,182). Considering the presence of multiple promoter sequences per gene, one representative promoter with the highest count per gene was selected, leaving n=20,094 unique gene promoters to calculate the frequencies. (*I*) Gene ontology (GO) analysis with Metascape for genes annotated by HOMER to promoter regions with ≥2 ZBED2 motif counts. Terms are ranked by their significance (*p* value) and the most significant term is highlighted. (*J*) ChIP-seq profiles of FLAG-ZBED2 in AsPC1 cells and H3K27ac in HPAFII cells at the promoters of the interferon genes *MX1*, *CMPK2* and *STAT2*. See also Fig. S3.

Since other ZBED zinc fingers function as sequence specific DNA binding domains (30, 31), we attempted to define a ZBED2 motif that correlates with its genomic occupancy. Using a *de novo* motif discovery analysis in the MEME-ChIP software (45), we derived an 8-nucleotide position weight matrix that closely correlated with ZBED2 enrichment observed by ChIP-seq (Fig. 3*E*). In addition, we found that the presence of two or more ZBED2 motifs correlated with strong genomic occupancy as well as with a stronger repressive effect on transcription than peaks displaying only a single ZBED2 motif (Fig. 3*F* and *G* and *SI Appendix*, Fig. S3*B*). Notably, we found that less than 1% of all human promoters of protein coding genes contain two or more ZBED2 motifs, and these promoters are strongly enriched for those driving expression of genes in the IFN pathway (Fig. 3*H* and *I* and Dataset S8). *MX1*, *CMPK2* and *STAT2* are examples of ISGs that contain two ZBED2 motifs in their promoters (Fig. 3*J*). Collectively, these results suggest that ZBED2 functions as a sequence-specific DNA-binding repressor of select ISG promoters harboring multiple ZBED2 motifs.

### Antagonistic regulation of ISG promoters by ZBED2 and IRF1

In our MEME analysis of ZBED2-enriched locations in the human genome, we noticed a strong correlation between ZBED2 occupancy and a motif recognized by the IRF family TFs (Fig. 4*A* and *SI Appendix*, Fig. S4*A*). RNA-seq analysis revealed variable expression of IRF family TFs in PDA cell lines, however the levels of *ZBED2* expression were closely correlated with that of *IRF1* (Fig. 4*B* and *SI Appendix*, Fig. S4*B* and *C*). A significant positive correlation between *ZBED2* and *IRF1* expression was also observed across all 1,156 cell lines within the CCLE database (*SI Appendix*, Fig. S4*D*). We next performed ChIP-seq analysis of IRF1 in AsPC1 cells, which confirmed its overlap with ZBED2 occupancy (*SI Appendix*, Fig. S4*E* and Dataset S9). For further analysis, we defined a high-confidence set of IRF1/ZBED2 co-occupied sites (Fig. 4*C* and Dataset S10). Notably the presence of IRF1 and ZBED2 co-occupancy was a feature that was highly enriched for ISG promoters, such as *CMPK2* and *STAT2* (Fig. 4*D* and *E* and *SI Appendix*, Fig. S4*F-I*).

**Fig. 4.**
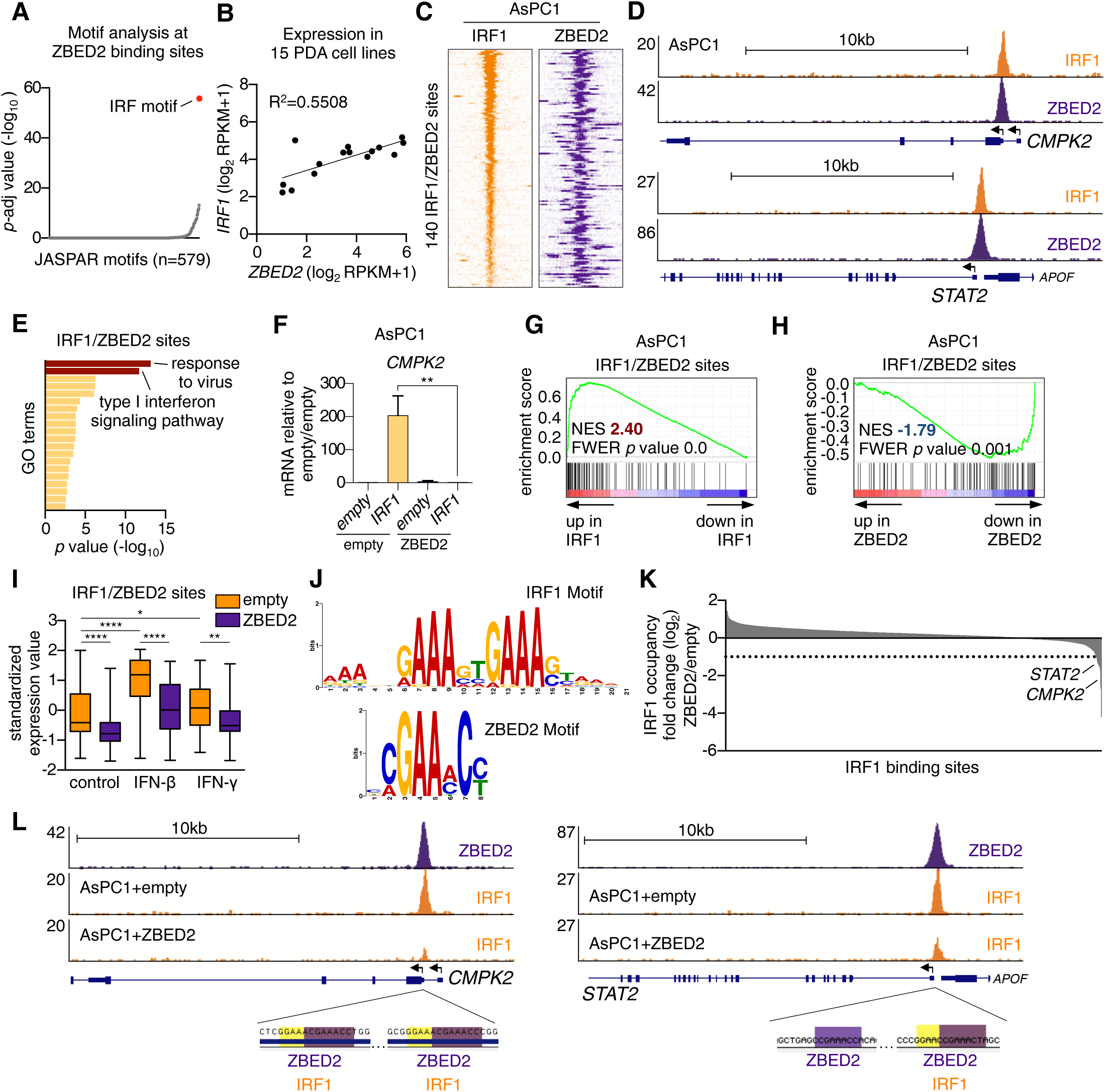
Antagonistic regulation of ISG promoters by ZBED2 and IRF1. (*A*) CentriMo motif enrichment analysis for JASPAR motifs at ZBED2 binding sites. (*B*) *ZBED2* expression versus the *IRF1* expression across 15 human PDA cell lines. (*C*) Density plot showing IRF1 and FLAG-ZBED2 enrichment surrounding a 2-kb interval centered on the summit of 140 intersecting IRF1 and FLAG-ZBED2 sites in AsPC1 cells, ranked by IRF1 peak intensity. (*D*) ChIP-seq profiles of IRF1 and FLAG-ZBED2 in AsPC1 cells at the promoters of *CMPK2* and *STAT2*. (*E*) Gene ontology (GO) analysis with Metascape of genes annotated by HOMER to IRF1/ZBED2 sites. Terms are ranked by their significance (*p* value) and the most significant terms (-log_10_ *p* value >12) are shown. (*F*) RT-qPCR analysis of *CMPK2* in AsPC1-empty and AsPC1-ZBED2 cells following IRF1 cDNA expression. Mean+SEM is shown. n=3. **p <0.01 by Student’s t test. (*G* and *H*) GSEA plots evaluating protein coding genes annotated by HOMER to IRF1/ZBED2 sites upon IRF1 (*G*) or ZBED2 (*H*) cDNA expression in AsPC1 cells. (*I*) Expression levels of protein coding genes annotated to IRF1/ZBED2 sites following 12-hour treatment with 0.2ng/µl of IFN-β, IFN-γ or control. \*\*\*\**p* <0.0001, \*\*\**p*<0.001, \*\**p*<0.01 *p <0.05 by one-way ANOVA. (*J*) IRF1 motif logo from the JASPAR database (top panel) and the ZBED2 motif logo (bottom panel). (*K*) ChIP-seq analysis showing the log_2_ fold change in IRF1 occupancy at IRF1 binding sites in AsPC1-ZBED2 versus AsPC1-empty cells. (*L*) ChIP-seq profiles of FLAG-ZBED2 and IRF1 at the promoters of *CMPK2* (left panel) and *STAT2* (right panel) in AsPC1-empty or AsPC1-ZBED2 cells. ZBED2 and IRF1 motifs recovered by FIMO (*p*<0.001) are highlighted in purple and yellow, respectively. See also Fig. S4.

Since ZBED2 is a transcriptional repressor and IRF1 is a transcriptional activator (Fig. 3*D*), we reasoned that these two TFs might function in an antagonistic manner to regulate ISG expression. Consistent with this hypothesis, RT-qPCR analysis showed that IRF1 led to potent activation of *CMPK2* while this effect was ablated in the presence of ZBED2 (Fig. 4*F*). RNA-seq analysis further verified that IRF1 and ZBED2 function in an antagonistic manner to regulate the entire program of co-occupied sites (Fig. 4*G* and *H* and *SI Appendix*, Fig. S4*J-M*). Considering these observations, we next treated AsPC1-empty and AsPC1-ZBED2 cells with IFN-β, IFN-γ or control for 12 hours and performed RNA-seq analysis. This analysis revealed that genes associated with IRF1/ZBED2 co-occupied sites were significantly induced by IFN and this effect was attenuated by the presence of ZBED2 (Fig. 4*I* and *SI Appendix*, Fig. S4*N*). Taken together, these data indicate that ZBED2 co-localizes with IRF1 at a subset of ISG promoters that are potently modulated by IFN pathway activation.

We noticed that the IRF1 motif and ZBED2 motif bear similarity with one another, with both factors able to recognize a GAAA sequence (Fig. 4*J*) (46). This led us to hypothesize that IRF1 and ZBED2 might bind to specific promoter sequences in a competitive manner, and this may contribute to the antagonistic function of these two TFs. To evaluate this, we performed IRF1 ChIP-seq in AsPC1 cells that either express or lack ZBED2. This analysis revealed a two-fold reduction of IRF1 occupancy at ∼19% of IRF1/ZBED2 co-bound sites (Fig. 4*K* and Dataset S11). This included sites at the promoters of ISGs *STAT2* and *CMPK2* (Fig. 4*K* and *L*), which were consistently observed to be among the most strongly downregulated genes upon ZBED2 expression (Fig. 2*E* and 3*C*). Notably, these two promoters both possess motifs with a common GAAA core that are capable of being recognized by both IRF1 and ZBED2 (Fig. 4*L*). Taken together, these findings reveal two distinct mechanisms by which ZBED2 can block the output of IRF1 at ISG promoters: through an intrinsic repressor function and via competitive DNA binding at specific motifs.

### ZBED2 protects PDA cells from IRF1- and interferon-γ-induced growth arrest

In addition to coordinating an anti-viral cellular response, IRF1 leads to a powerful growth arrest when activated in tumor cells (47–49). Consistent with this observation, we found that forced IRF1 expression in several human and murine PDA cell line contexts resulted in a significant growth arrest (Fig. 5*A* and *SI Appendix*, Fig. S5*A*). Remarkably, if we repeated this experiment in the presence of ZBED2, the IRF1-induced growth arrest was prevented (Fig. 5*B-E*). To verify this result in the setting of a more physiological context, we used exogenous IFN-γ to promote IRF1 activation in PDA cells. Notably, in PDA cell lines, IRF1 knockout only blocked IFN-γ-induced growth arrest but not growth arrest caused by IFN-β, which instead relied on IRF9 (Fig. 5*F-H* and *SI Appendix*, Fig. S5*B-D*). Given that the growth inhibition effects of IFN-γ stimulation are at least partially mediated through IRF1 induction, we reasoned that ZBED2 expression might confer a protective effect in this context. Consistent with this hypothesis, ZBED2 expression was sufficient to protect PDA cells from the effects of IFN-γ-mediated growth arrest, which was observed in AsPC1 cells, as well as in three murine PDA cell lines derived from the KPC (*Kras^+/LSL-G12D^*; *Trp53^+/LSL-R172H^*; *Pdx1-Cre*) mouse model (Fig. 5*I* and *J* and *SI Appendix*, Fig. S5*E*) (50, 51). Taken together, these data suggest that ZBED2 can antagonize both the transcriptional and phenotypic consequences of IRF1 activation in PDA cells.

**Fig. 5.**
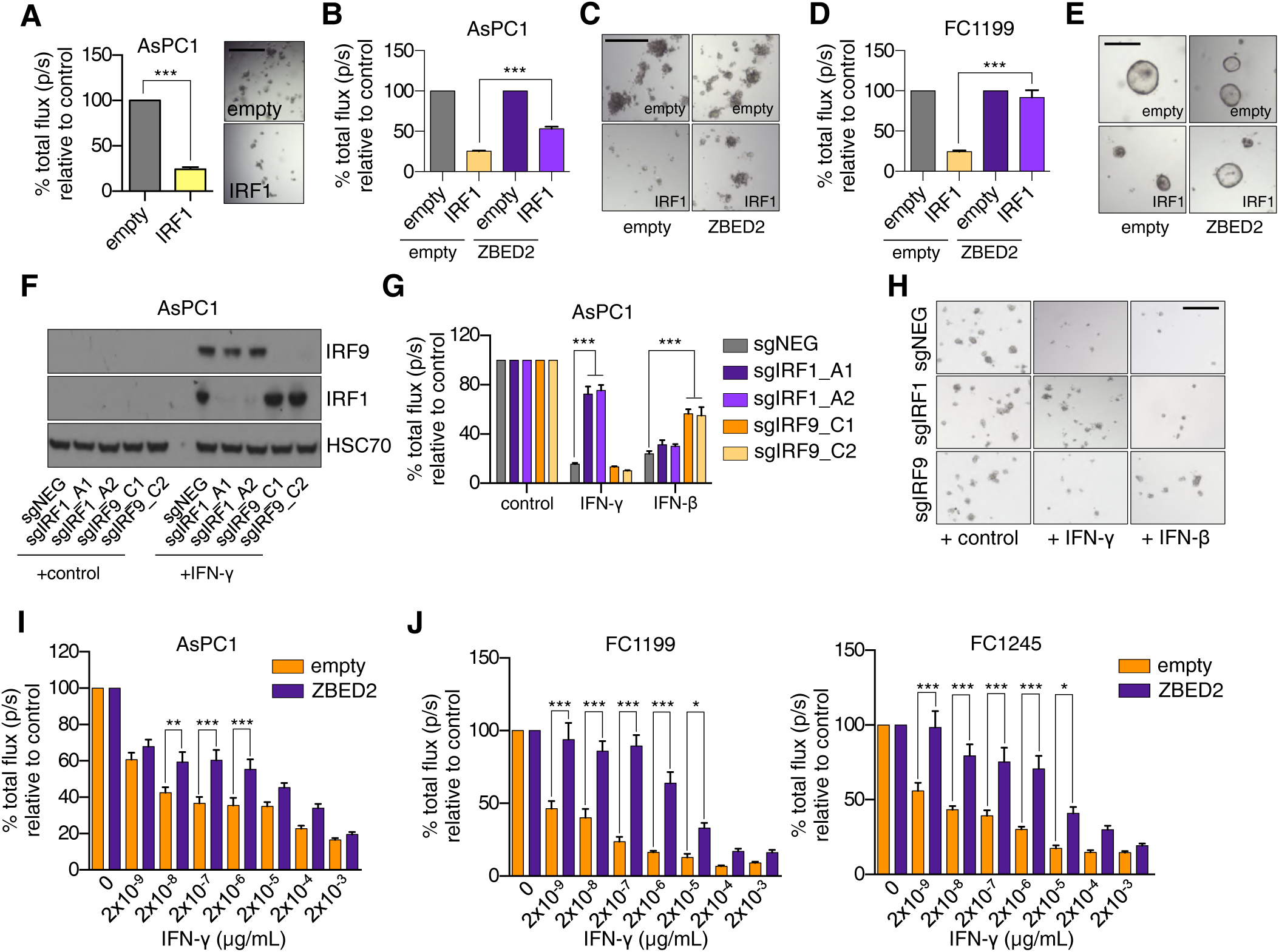
ZBED2 protects PDA cells from IRF1- and interferon- γ -induced growth arrest. (*A*) Luciferase-based quantification of cell viability of AsPC1 cells grown in Matrigel on day 7 post infection with IRF1 cDNA or the empty vector. Representative bright field images (right panel) are shown. Scale bar indicates 200µm. (*B*) Luciferase-based quantification of cell viability of AsPC1-empty and AsPC1-ZBED2 cells grown in Matrigel following co-expression of IRF1 cDNA or empty vector for 7 days. (*C*) Representative bright filed images from (*B*). Scale bar indicates 500µm. (*D*) Luciferase-based quantification of cell viability of KPC-derived FC1199 cells stably expressing ZBED2 or the empty vector grown in Matrigel following co-expression of IRF1 cDNA or empty vector for 7 days. (*E*) Representative bright filed images from (*D*). Scale bar indicates 200µm. (*F*-*H*) AsPC1 cells infected with sgRNAs targeting IRF1, IRF9 or a control sgRNA (sgNEG) were plated in Matrigel and grown for 7 days in the presence of 20ng/µl of IFN-γ, IFN-β or control. Representative western blots following overnight stimulation with 20ng/µl of IFN-γ (*F*), luciferase-based quantification (*G*) and representative bright field images on day 7 of the assay (*H*) are shown. Scale bar indicates 500µm. (*I* and *J*) AsPC1 cells (*I*) or the indicated mouse KPC cell lines (*J*) were infected with the ZBED2 cDNA or an empty vector and grown in Matrigel with the increasing concentrations of IFN-γ. Bar charts show luciferase-based quantification on day 7. Mean+SEM is shown. n=3. For (*A*), (*B*), (*D*) and (*G*), ***p <0.001 by Student’s t test. For (*I*) and (*J*), ***p<0.001, **p<0.01 *p <0.05 by one-way ANOVA. See also Fig. S5.

### ZBED2 represses pancreatic progenitor lineage identity in PDA

Prior transcriptome analyses of tumor samples have revealed two major molecular subtypes of PDA: a ‘pancreatic progenitor’ and a ‘squamous’ subtype (13, 35, 36, 52). The pancreatic progenitor subtype is also known as classical PDA, and expresses endodermal TFs (e.g. GATA6, HNF4, FOXA) at high levels. In contrast, the squamous subtype (also known as basal-like) silences the pancreatic progenitor gene signature and instead expresses markers of the squamous lineage (e.g. KRT5, TP63) in association with inferior patient survival outcomes (10, 13). Remarkably, we found in three independent human tumor transcriptome datasets (13, 35, 36), *ZBED2* expression was highly associated with squamous subtype PDA tumors (Fig. 6*A-F* and Dataset S12). This prompted us to investigate whether ZBED2 has causal role influencing the ‘pancreatic progenitor’ or ‘squamous’ transcriptional signatures in PDA cell lines. Using the aforementioned RNA-seq analysis, we found that the pancreatic progenitor signature was significantly repressed by ZBED2 expression in 13/15 PDA cell lines examined (Fig. 6*G* and *H*). The two exceptions were MIAPaca2 and PANC1 cells, which do not express pancreatic progenitor signature genes (10). The squamous signature was more variably affected by ZBED2 expression (*SI Appendix*, Fig. S6*A*). In accord with previous findings (27), we found that IRF1 activates the pancreatic progenitor signature, which is consistent with its antagonism with ZBED2 (Fig. 6*I*). We noticed that one of the IRF1/ZBED2 co-occupied sites was found at the promoter of the *GATA6* gene, which encodes an established marker of the pancreatic progenitor transcriptional program (Fig. 6*J* and *K*) (36, 52–54). Consistent with their antagonistic activities, we found ZBED2 or IRF1 expression resulted in the repression or activation of *GATA6* in this context, respectively (Fig. 6*L* and *SI Appendix*, Fig. S6*B* and *C*). Of note, out of the 15 PDA cell lines used in this study, we observed the lowest levels of *GATA6* and *IRF1* in MIAPaca2 and PANC1 cells (*SI Appendix*, Fig. S6*D* and *E*).

**Fig. 6.**
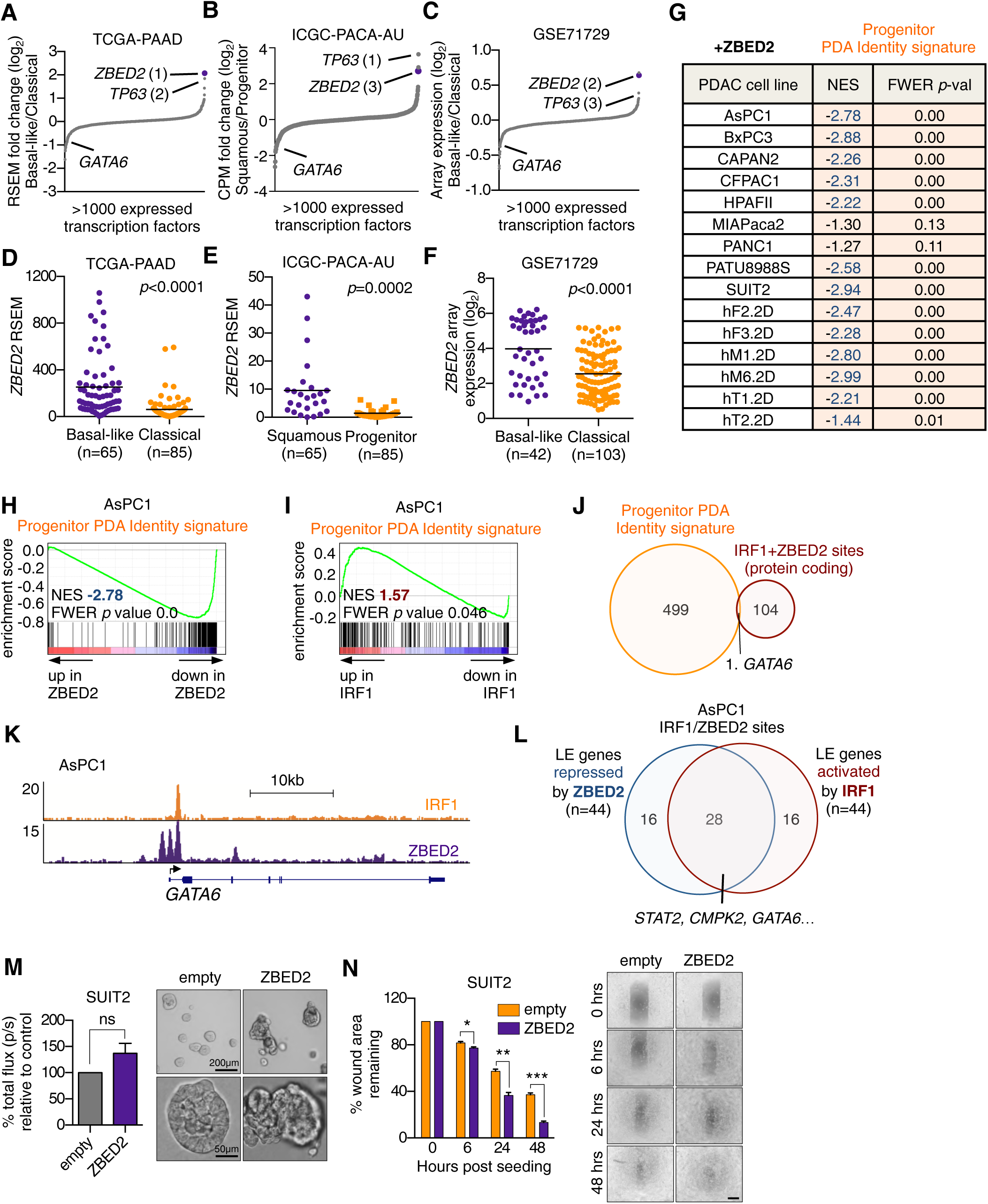
ZBED2 represses pancreatic progenitor lineage identity in PDA. (*A*-*C*) TF expression in Basal-like/Squamous and Classical/Progenitor subtypes of PDA. TFs are ranked by their mean log_2_ fold change in Basal-like versus Classical (*A* and *C*) or Squamous versus Progenitor (*B*) patient samples from the indicated studies. (*D*-*F*) *ZBED2* expression in PDA patient samples stratified according to molecular subtype. Each dot represents one patient sample. *p* value was calculated using Student’s t test. (*G*) Summary of GSEA evaluating the Progenitor PDA Identity signature upon ZBED2 cDNA expression in 15 PDA cell lines. (*H* and *I*) GSEA plots evaluating the Progenitor-PDA Identity signature following expression of ZBED2 (*H*) or IRF1 (*I*) in AsPC1 cells. (*J*) Overlap of Progenitor PDA Identity genes with protein coding genes associated with IRF1/ZBED2 sites. (*K*) ChIP-seq profiles of IRF1 and FLAG-ZBED2 at the promoter of *GATA6* in AsPC1 cells. (*L*) Overlap of leading edge (LE) genes associated with IRF1/ZBED2 sites that are repressed by ZBED2 or activated by IRF1 in AsPC1 cells. (*M*) Quantification of colony size in three-dimensional (3D) Matrigel colony formation assays. Means+SEM are shown. n=3. Representative images at day 7 are shown (right panel). (*N*) Quantification of scratch assays at the indicated time points post-seeding and representative images (right panel). Scale bar indicates 500µm. Means+SEM are shown. n=3. ***p<0.001, **p<0.01 *p <0.05 by Student’s t test. See also Fig. S6.

Consistent with the results above, patient tumor samples with high levels of *ZBED2* expression were much more likely to present with poorly differentiated tumors compared to those patients with low *ZBED2* expression (*SI Appendix*, Fig. S6*F* and *G*). Moreover, we found that ZBED2 expression in PDA cells was sufficient to promote loss of epithelial integrity and increased invasiveness when cells were plated in Matrigel (Fig. 6*M*), as well as enhanced migratory behavior in scratch assays (Fig. 6*N*). These findings are in support of ZBED2 regulating the lineage identity of PDA cells and can account for the biased expression of *ZBED2* in squamous subtype PDA tumors.

## DISCUSSION

Here, we have shown that a previously uncharacterized zinc finger protein ZBED2 is aberrantly expressed in a diverse collection of human tumors in a manner that correlates with poor clinical outcomes. Using PDA cells as an experimental system, we have defined two major molecular functions of ZBED2 that may underpin its role in promoting cancer: as an antagonist of the IFN response and as a modulator of epithelial cell identity. In addition, we provide evidence that both of these functions of ZBED2 can be accounted for by antagonism of IRF1-mediated transcriptional activation.

A key mechanistic advance in our study is in revealing the DNA sequence motif recognized by ZBED2, which we show is disproportionately enriched at ISG promoters in juxtaposition with IRF1 motifs. This observation suggests that the evolved function of ZBED2 in mammalian biology is as a tissue-specific attenuator of the IFN response. The tissues that express *ZBED2* at the highest levels include thyroid, esophagus, lung, and skin, which are organs that are often exposed to pathogens and may employ ZBED2 to define thresholds for IFN responsiveness. Notably, inflammation of the thyroid is one of the most common side effects of interferon-α therapy in humans (55), which leads us to speculate that ZBED2 may have evolved to protect vulnerable tissues from the damaging effects of chronic IFN stimulation. Other ZBED TFs have also been implicated in the regulation of immune responses. For example, ZBED1 has been described as a viral restriction factor that negatively regulates viral growth (56). In addition, a recent study in plants identified a ZBED protein as a novel regulator of immunity in this context (57). Thus, the function of ZBED2 defined in our study may have its origins in the early evolutionary functions of ZBED proteins as important regulators of immune responses.

We have defined ZBED2 as an attenuator of IFN responses through its ability to antagonize the transcriptional output of IRF1. Importantly, IRF1 is a validated tumor suppressor (47–49, 58) and IFN pathways have a well-established role in the regulation of antitumor immunity (59). Thus, antagonism of IRF1 provides a plausible explanation for why high ZBED2 expression is selected for during tumorigenesis. Less clear are the initial triggers of IRF1/IFN pathway activation in PDA. A recent study demonstrated that cancer cells exhibit high ISG expression due to chronic IFN signaling sustained by the STING adaptor protein, which is activated by various intracellular nucleic acid sensors (60, 61). Thus, elevated ISG expression in cancer cells could be attributed to the accumulation of cytosolic double-stranded DNA, which may arise from the disruption of micronuclei as a result of DNA damage, chromosome mis-segregation, and chromothripsis (62, 63). Such features of high genomic instability have been strongly implicated in the development and progression of PDA (43, 64). However, sustained activation of ISG expression via the STING pathway was shown to be dependent on chronic production of type I IFNs (60). In our analysis of human PDA cell lines, we observed high basal levels of ISG expression in PDA cells in a manner that is independent of type I IFNs. While the triggers of the ISG pathway in PDA await further characterization, our results strongly suggest that the function of ZBED2 in this context is to block the anti-tumor effects of IRF1 within the IFN pathway.

Histopathological and transcriptome studies have shown that a subset of aggressive PDA tumors undergo trans-differentiation into the squamous lineage (13, 65–67). In addition to the aberrant upregulation of squamous lineage markers, a key attribute of these tumors is the silencing of genes associated with pancreatic progenitor cell identity. A key finding in our study is that ZBED2 is selectively upregulated in squamous-subtype PDA tumors, and in this context represses the pancreatic progenitor transcriptional program. Notably, we also observe high *ZBED2* expression in normal squamous epithelial tissues, such as the skin, lung and esophageal mucosa, suggesting a normal function for ZBED2 in the regulation of squamous epithelial cells. Interestingly, single-cell RNA-seq analysis of human foreskin keratinocytes recently identified ZBED2 as a key TF in the regulation of keratinocyte differentiation programs (33). ZBED2 was predicted to promote the basal keratinocyte state and experimental depletion of ZBED2 was shown to induce keratinocyte differentiation, demonstrated by the upregulation of *KRT10* (33). It is noteworthy that IRF family members have also been ascribed functional roles in regulation of the skin epidermis. For example, both IRF2 and IRF6 are necessary for regulating proliferation and terminal differentiation of keratinocytes within the epidermis (68–71). Thus, it is possible that the ability of ZBED2 to modulate the transcriptional output of IRF TFs may extend to its ability to regulate stem cell function and cell identity within normal squamous epithelial tissues.

Similar to ZBED2, prior studies have implicated TP63 and GLI2 as additional determinants of squamous cell identity in PDA (10, 11, 13, 15). Our analysis has shown that ZBED2 and TP63 are among the most aberrantly expressed TFs in the squamous subtype of PDA, however these two TFs are not expressed in a mutually exclusive manner. These findings suggest the aberrant expression of TP63, GLI2, or ZBED2 offer distinct mechanisms by which pancreatic cancer cells with a ductal identity adopt squamous features during the pathogenesis and progression of this disease. An important area of future investigation will be to determine whether such factors can function in a collaborative manner to regulate squamous identity in this disease.

An important feature of squamous subtype tumors is that they display poorly differentiated histopathological features and an exceptionally poor prognosis (13, 35, 36, 52, 72, 73). Interestingly, a recent study identified IRF1 as one of several candidate TFs responsible for preserving differentiation and epithelial identity in human PDA (27). In this study, IRF1 was shown to associate with ‘low-grade’ enhancers controlling grade-specific transcriptional programs, which is in accord with our functional studies of IRF1. We have shown that a key IRF1/ZBED2 co-occupied target gene is *GATA6*, which is overexpressed and amplified in the pancreatic progenitor subtype of PDA (52, 54, 74). Indeed, *GATA6* expression is altered by multiple mechanisms in pancreatic tumors and was found to be a robust biomarker for PDA subtypes (36, 73). Thus, our study further reinforces how cell identity in PDA is defined by antagonism among TFs important for maintaining the epithelial identity of this lineage.

## MATERIALS AND METHODS

### Cell Lines and Cell Culture

Human PDA cell lines were cultured in RPMI 1640 (Gibco) containing 10% FBS. Mouse PDA cell lines, HEK 293T cells and Plat-E cells were cultured in DMEM containing 10% FBS. Plat-E and HEK 293T cells were used for packing retrovirus and lentivirus, respectively. All cells were cultured with 1% L-glutamine and 1% Penicillin/Streptomycin at 37°C with 5% CO_2_. All cell lines were routinely tested for mycoplasma. Interferon treatments of cell lines were performed with IFN-β (PeproTech; #300-02BC), human IFN-γ (PeproTech; #300-02), or mouse IFN-γ (PeproTech; #315-05).

### Plasmid Construction

ZBED2 and IRF1 cDNA was PCR amplified from oligo-dT primed cDNA from gDNA harvested from CAPAN2 cells, gel purified and cloned into LentiV_Puro vector (addgene #111886) using In-Fusion following the manufacturer’s instructions (Clontech: #638909). A 3*FLAG sequence was added to the N-terminus of ZBED2 cDNA to generate the FLAG-ZBED2 construct.

### Lentiviral Production and Infection

Lentivirus was produced in HEK 293T cells by transfecting plasmid DNA and packaging plasmids (VSVG and psPAX2) using Polyethylenimine (PEI 25000; Polysciences; Cat# 23966-1). Media was replaced with target media 6-8 hours following transfection and lentivirus-containing supernatant was subsequently collected every 12 hours for 48 hours prior to filtration through a 0.45µm filter. For infection of human and mouse PDA cells, cell suspensions were mixed with lentiviral-containing supernatant supplemented with polybrene to a final concentration of 4µg/ml. Cells were plated in tissue culture plates of the appropriate size and lentiviral-containing supernatant was replaced with fresh media after an incubation period of 24 hours. At 48 hours post-infection, transduced cells were selected with puromycin (3µg/ml) or G418 (1mg/ml) for five days prior to RNA extraction, lysate preparation and/or phenotypic analyses on day seven post infection.

### Retroviral Production and Infection

Retrovirus was produced in Plat-E cells by transient transfection with PEI (Polysciences; Cat#23966-1) according to standard protocols. The viral supernatants were harvested at 48 and 72 hours post-transfection prior to filtration through a 0.45µm filter. To maximize transduction efficiency, PANC0403 cells were infected with retrovirus by centrifugation at 2,000 rpm for 30 mins in the presence of 8µg/ml polybrene. Retroviral-containing supernatant was replaced with fresh media after an incubation period of 24 hours. At 48 hours post-infection, transduced cells were selected with G418 (1mg/ml) for five days prior to RNA extraction.

### CRISPR-based Targeting

To generate cell lines in which ZBED2, IRF1, IRF9 or IFNAR1 had been stably knocked out, PDA cells expressing Cas9 in the LentiV-Cas9-puro vector (addgene # 108100) were infected in a pooled fashion with domain-targeting sgRNAs or a control sgRNA (sgNEG) in the LRG2.1_Neo vector (addgene #125593). sgRNA targeting sequences can be found in Dataset S13.

### shRNA-based Targeting

shRNAs targeting ZBED2 or control were cloned into the miR30-based retroviral shRNA expression vector LENC (LTR-miR30-shRNA-PGK-neo-mCherry) (#111163) (75). Two days post infection with shRNAs, transduced cells were selected with 1mg/ml of G418 for five days prior to RNA extraction. shRNA sequences can be found in Dataset S13.

### RNA extraction and RT-qPCR analysis

Total RNA was extracted using TRIzol reagent following the manufacturer’s instructions. 1µg of total RNA was reverse transcribed using qScript cDNA SuperMix (Quanta bio; 95048-500), followed by RT-qPCR analysis with Power SYBR Green Master Mix (Thermo Fisher Scientific; 4368577) on an ABI 7900HT fast real-time PCR system. Gene expression was normalized to *GAPDH*. RT-qPCR primers used can be found in Dataset S13.

### Cell Lysate Preparation and Western Blot Analysis

Cell cultures were collected and 1×10^6^ cells were counted by trypan blue exclusion, washed with ice cold PBS, resuspended in 100µl PBS and lysed with 100µl of 2x Laemmli Sample Buffer supplemented with β-mercaptoethanol by boiling for 30 minutes. Samples were centrifuged at 4°C for 15 mins at 10,000xg and the supernatant was used for western blot analysis with standard SDS-PAGE-based procedures. Primary antibodies used were ZBED2 (Sigma-Aldrich; SAB1409809), IRF1 (Cell Signaling; #8478), IRF9 (Cell Signaling; #76684), MX1 (Cell Signaling; #37849), pSTAT2 (Cell Signaling; #88410), pSTAT1 (Cell Signaling; #9167), STAT2 (Cell Signaling; #72604), CMPK2 (abcam; ab103658), HSC70 (Santa Cruz; sc-7298) and ACTB (Sigma-Aldrich; A3854). Proteins were detected using HRP-conjugated secondary antibodies.

### GAL4 Fusion Reporter Assays

To measure transcriptional activation activity of ZBED2 and IRF1, the GAL4 DNA-binding domain was fused to full length ZBED2 and IRF1 cDNA and these constructs were individually co-transfected into HEK 293T cells with the pGL4.35[luc2P/9XGAL4UAS/Hygro] vector (Promega, #E1370). 48 hours post transfection, an equal volume of Dual-Glo luciferase buffer and Dual-Glo Stop&Glo buffer were added sequentially and relative luciferase activity was measured according to manufacturer’s instructions. Firefly luciferase activity was normalized to internal Renilla luciferase activity.

### In Vitro Luciferase Imaging and 3D Matrigel Assays

To generate luciferase expressing PDA cell lines, cells were infected with a luciferase transgene in a Lenti-luciferase-blast vector (10) and stable cell lines were generated by selection with 10µg/ml blasticidin. Following CRISPR-based targeting or cDNA expression, cells were plated at a density of 5×10^3^−2×10^4^ cells per well of a black, clear-bottom, ultra-low attachment 96-well plate (10014-318; CELLSTAR) in 200µl of RPMI supplemented with 2% FBS and 5% Matrigel with or without the addition of human IFN-γ (PeproTech; #300-02), mouse IFN-γ (PeproTech; #315-05) or control. Cells were imaged on day seven post-plating using an IVIS Spectrum system (Caliper Life Sciences) six minutes post addition of D-Luciferin (150µg/ml) to each well. Bright field images were also captured on day seven post plating.

### Scratch Assays

Cells were first plated to confluency in triplicate in wells of a standard 24-well plate. At day 0 of the assay, a wound was applied down the center of the well using a pipette tip. Media was subsequently removed and cells washed with PBS before addition of 1ml serum-free RPMI. Bright field images were captured using a 4x objective immediately (0 hours) and then at 6 hours, 24 hours and 48 hours post-plating. Area of the wound was quantified using ImageJ software (NIH).

### RNA-seq Library Construction

RNA-seq libraries were constructed using the TruSeq sample Prep Kit V2 (Illumina) according to the manufacturer’s instructions. Briefly, 2µg of purified RNA was poly-A selected and fragmented with fragmentation enzyme. cDNA was synthesized with Super Script II Reverse Transcriptase (Thermo Fisher; 18064014), followed by end repair, A-tailing and PCR amplification. RNA-seq libraries were single-end sequenced for 50bp using an Illumina NextSeq platform (Cold Spring Harbor Genome Center, Woodbury).

### RNA-Seq Data Analysis

Single end 50bp sequencing reads were mapped to the hg19 or hg38 genomes using HISAT2 with standard parameters (76). Structural RNA was masked and differentially expressed genes were identified using Cuffdiff (77). All the following analysis was performed on genes with an RPKM value no less than 2 in either control or experimental samples. To generate ranked gene lists for Pre-ranked GSEA, genes were ranked by their mean log_2_ fold change between the two experimental groups of interest. For unbiased interrogation of the transcriptional consequences following ZBED2 knockout in PANC0403 cells or ZBED2 cDNA expression in HPAFII cells, ranked gene lists were ran against the H1, C2, C5 and C6 gene sets within the MSigDB v7.0 (n = 15,736) with those with ≤15 or ≥500 expressed genes filtered out, leaving 7,573 and 7,593 interrogated genes sets for PANC0403 and HPAFII cells, respectively. The Interferon Response gene signature was defined by merging the Broad Institute’s Hallmark Interferon Alpha Response and Interferon Gamma Response gene signatures to generate a gene set comprising 222 unique gene symbols. The Progenitor-PDA and Squamous-PDA Identity signatures have been previously described (10). Heat maps of expression values were generated using Morpheus from the Broad Institute (software.broadinstitute.org/morpheus). Gene ontology analysis was performed using Metascape (78).

### ChIP-Seq Library Construction

ChIP procedures were performed as previously described (10). Antibodies used for ChIP-seq in this study were IRF1 (Cell Signaling; #8478) and FLAG (Sigma-Aldrich; F1804). ChIP-seq libraries were constructed using Illumina TruSeq ChIP Sample Prep kit following manufacture’s protocol. Briefly, ChIP DNA was end repaired, followed by A-tailing and size selection (300-500bp) by gel electrophoresis using a 2% gel. 15 PCR cycles were used for final library amplification that was analyzed on a Bioanalyzer using a high sensitivity DNA chip (Agilent). ChIP-seq libraries were single-end sequenced for 50bp using an Illumina NextSeq platform (Cold Spring Harbor Genome Center, Woodbury).

### ChIP-Seq Data Analysis

Single end 50bp sequencing reads were mapped to the hg19 genome using Bowtie2 with default settings (79). After removing duplicated mapped reads using SAM tools (80), MACS 1.4.2 was used to call peaks using input genomic DNA as control (81). Only peaks enriched greater than or equal to 10-fold over input samples were used for subsequent analyses. FLAG-ZBED2 peaks that were >10 fold over input with >5 tags per million in both the AsPC1 and SUIT2 datasets (overlapping intervals by at least 1bp) were taken forward as high confidence ZBED2 peaks (n = 2,451). De novo motif discovery was performed on DNA sequences extending 200bp upstream and downstream of the top 1,000 high confidence peak summits using MEME-ChIP (45) from the MEME Suite (82). The same DNA sequences were used to identify motifs enriched at ZBED2 binding sites using CentriMo via the MEME Suite with default settings (83). For this analysis, the non-redundant JASPAR core vertebrate database (v2018) containing 579 transcription factor motifs were interrogated. Annotation of ChIP-seq peaks was performed using HOMER v4.9 with default settings (84). To interrogate expression changes of ZBED2 bound genes in HPAFII cells, the top 500 unique, protein coding genes that were expressed and annotated to ZBED2 peaks (ranked by ZBED2 binding intensity) were defined as ZBED2 bound genes used for GSEA. To identify high confidence IRF1 peaks in AsPC1 cells, cells were first treated overnight with control or IFN-γ (0.2ng/µl; PeproTech; #300-02) prior to ChIP to increase IRF1 expression. Peaks that were >10 fold over input from both the control and IFN-γ treated datasets were merged giving 11,783 peaks. Tag counts were recounted and those peaks with >5 tags per million in either dataset were taken forward as high confidence IRF1 peaks (n = 1,946). Peaks that may be recognized by both ZBED2 and IRF1 were defined as those intervals in high confidence ZBED2 and IRF1 peaks that overlap by at least 1bp. TreeView software was used to generate the heat-map density plots and the contrast was adjusted proportionally to the total uniquely mapped reads for visual comparison across samples (85). H3K27ac ChIP-seq in AsPC1, SUIT2 and HPAFII cells were from a previous study (10).

### ZBED2 Motif Scanning

The ZBED2 motif output from *de novo* motif discovery analysis was used for this analysis. Genomic DNA and transcriptional start site annotation were downloaded from GENCODE: GRCh37 v19 for human (n=73,182). Promoters were defined as 500 bp sequences upstream and 100 bp downstream of annotated transcriptional start sites of protein coding transcripts. FIMO (86) was used to scan both strands of each promoter DNA sequence using a p-value cutoff of 10^-4^ and default settings using the MEME suite. For GSEA evaluating ZBED2 bound genes at promoter regions with ≥2 motif counts upon ZBED2 cDNA expression, this analysis was restricted to 139 protein coding genes with RPKM expression values ≥2 in at least 10/15 of the PDA cell lines interrogated; for comparative analysis, a random set of 139 protein coding genes meeting the same criteria with 0 or 1 ZBED2 motifs in their promoter was used.

### Analysis of Publicly Available Datasets

For analysis of TF expression in primary PDA tumors versus normal pancreas tissue from the study by Moffitt et al (GSE71729), as well as Basal-like versus Classical tumors, TFs (87) with a mRNA array value ≥2 in at least 10% of samples (n=252 samples) were considered as expressed (n=1,079 TFs). For analysis of TF expression in Basal-like versus Classical tumors from the TCGA-PAAD study, TFs with a log_2_ RSEM value ≥6.1 in at least 10% of samples (n=150 samples) were considered as expressed (n=1,110 TFs). For analysis of TF expression in Squamous versus Progenitor tumors from the study by Bailey et al (ICGC-PACA-AU), TFs with a CPM value ≥4 in at least 10% of samples (n=96 samples) were considered as expressed (n=1,052 TFs). Patient survival analysis across the TCGA dataset was performed using a publically available web tool (40). Patient survival data for PDA studies were obtained from the cBioPortal (TCGA-PAAD) (88) and ICGC Data Portal (PACA-AU) (13) and were downloaded in January 2018. For the TCGA-PAAD study, only the 150 confirmed PDA cases were used for the analyses in this study (36). In the two PDA studies, samples were designated as *ZBED2*^high^ or *ZBED2*^low^ based on z-score expression values >0.35 or <0, respectively. For analysis of *ZBED2* expression in the TCGA PanCancer Atlas studies (39) and the Broad Institute’s CCLE (41), data was downloaded from the CBioPortal (88) in October 2019. Cell lines in the CCLE database with a *ZBED2* log_2_(RPKM+1) expression value ≥1.5, which approximates to cell lines in 75^th^ percentile, were stratified as *ZBED2*^high^; those less than this value were stratified as *ZBED2*^low^. Analysis of genetic alterations in *ZBED2*^high^ versus *ZBED2*^low^ cell lines was performed using the CBioPortal and 127 significantly mutated genes identified from 12 tumor types were interrogated (89).

### Statistical Analysis

For graphical representation of data and statistical analysis, GraphPad Prism was used and statistical significance was evaluated by *p* value using Prism software as indicated in the figure legends. Data are presented as mean with SEM and n refers to the number of biological repeats. For Kaplan-Meier survival curves, the log rank (Mantel-Cox) test was used to estimate median overall survival and statistical significance.

### Data and Software Availability

ChIP-seq and RNA-seq data reported in this paper is available via the GEO: GSE141607 with reviewer token: gpatcqewtxihfgt. Previously deposited ChIP-seq data (10) can be found here GEO: GSE115463. PDA patient microarray data (35) can be found here GEO: GSE71729.

## Acknowledgements

The authors would like to thank the Cold Spring Harbor Cancer Center Support Grant (CCSG) shared resources: Bioinformatics Shared Resource and Next Generation Sequencing Core Facility. We would also like to thank Kenneth Olive and Carlos Maurer (Columbia University, NY) for critical discussions during manuscript preparation and Noah Dukler (CSHL, NY) for phylogenetic analysis of ZBED family members. T.D.D.S. was supported by a grant from the State of New York, contract no. C150158. C.R.V. was supported by Pershing Square Sohn Cancer Research Alliance, the Cold Spring Harbor Laboratory and Northwell Health Affiliation, the National Cancer Institute (NCI) 5P01CA013106-Project 4 and 1RO1CA229699, the Thompson Family Foundation, the Simons Foundation and a Career Development Award from the Pancreatic Cancer Action Network-American Association for Cancer Research (AACR) 16-20-25-VAKO.

## Disclosures

C.R.V. has received funding from Boehringer-Ingelheim and is an advisor to KSQ Therapeutics.

## Supplementary Figures

**Fig. S1.**
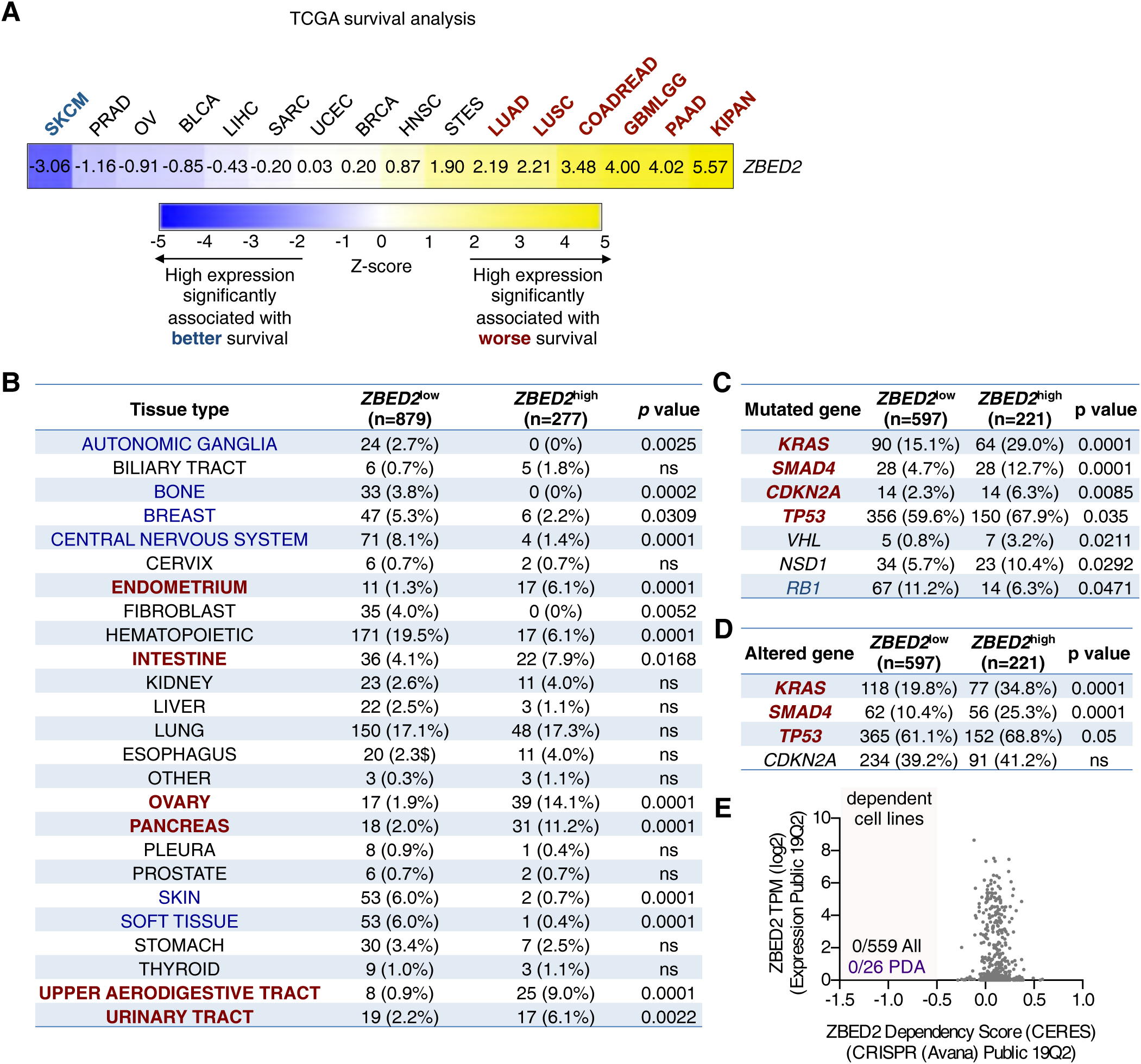
*ZBED2* is a poor prognosis marker in numerous cancers, is highly expressed in a subset of cancer cell lines and is dispensable for cell growth *in vitro*. Related to Fig. 1. (*A*) Survival analysis based on *ZBED2* expression levels across TCGA profiled tumors. Z-score values ≤-2 and ≥2 are considered significant. See Smith and Sheltzer (2018). KIPAN, pan kidney cancer; PAAD, pancreatic adenocarcinoma; GBMLGG, Glioblastoma/Glioma; COADREAD, colorectal adenocarcinoma; LUSC, Lung squamous cell carcinoma; LUAD, Lung adenocarcinoma; STES, Esophagus-Stomach Cancers; HNSC, Head and Neck squamous cell carcinoma; BRCA, Breast invasive carcinoma; UCEC, Uterine Corpus Endometrial Carcinoma; SARC, Sarcoma; LIHC, Liver hepatocellular carcinoma; BLCA, Bladder Urothelial Carcinoma; OV, Ovarian serous cystadenocarcinoma; PRAD, Prostate adenocarcinoma; SKCM, Skin Cutaneous Melanoma. (*B*) Proportion of cell lines from the CCLE stratified as *ZBED2*^low^ or *ZBED2*^high^ classified based on their tissue of origin. (*C* and *D*) Association of *ZBED2* expression with genetic features in CCLE cell lines. 127 significantly mutated genes in human cancer (Kandoth et al., 2013) were analyzed. For (*C*) only genetic mutations were considered and only significantly mutated genes are shown. For (*D*) both genetic mutation and copy number alteration (i.e. loss of a tumor suppressor or gain of an oncogene) were considered as altered genes. The table displays the top four most significantly altered genes in PDA. Statistical significance for the indicated comparisons was assessed using Fisher’s Exact Test, ns = not significant. (*E*) Summary of ZBED2 dependency score (CERES) and expression in 559 cancer cell lines from the depmap portal (depmap.org). The number of cell lines dependent on ZBED2 for cell growth across all cell lineages is indicated, as well as the 26 pancreatic cancer cell lines (PDA) included in this dataset. Cell lines with a CERES ≤-0.5 were considered as dependent.

**Fig. S2.**
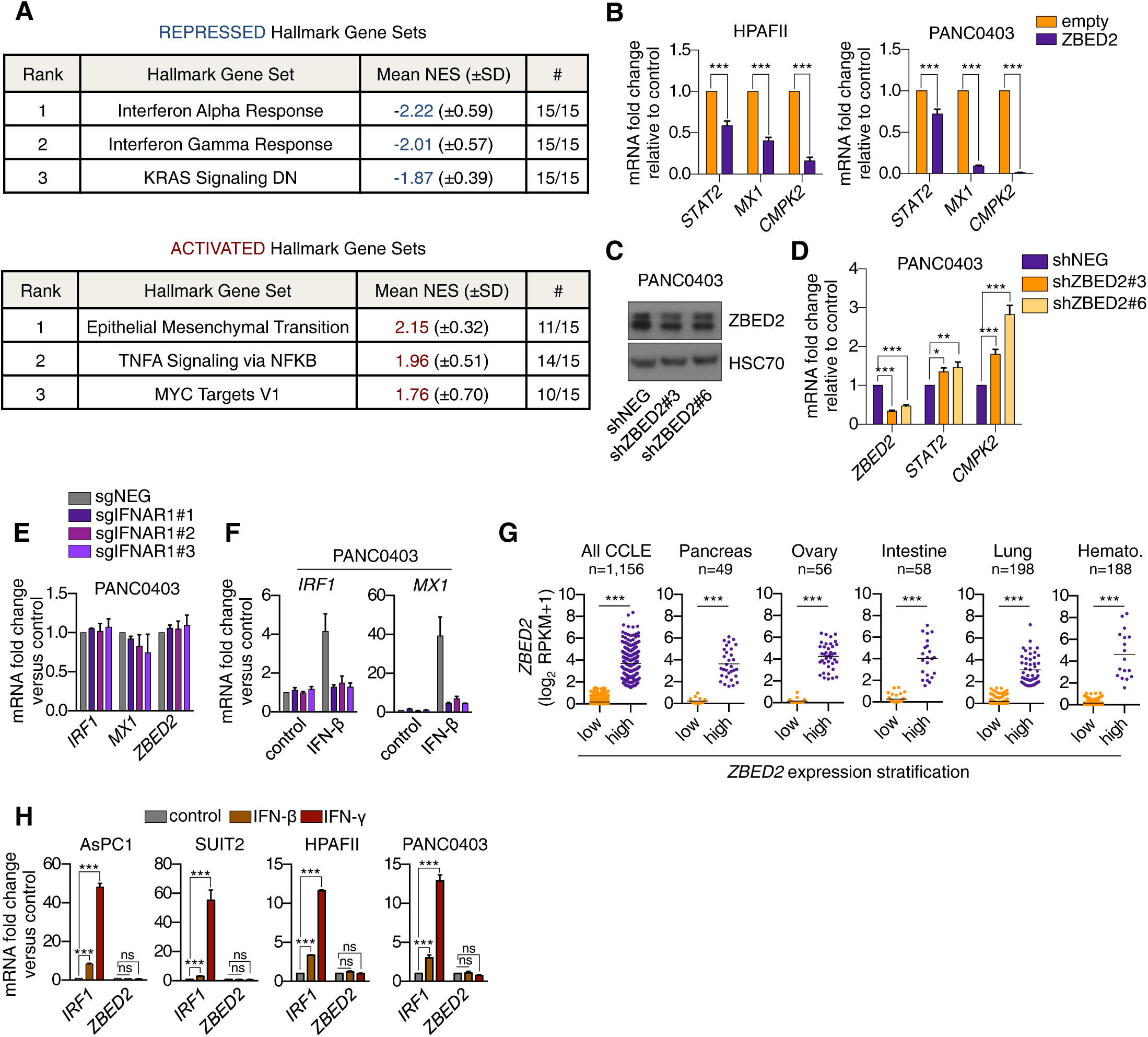
ZBED2 inhibits the expression of genes in the interferon response pathway. Related to Fig. 2. (*A*) Summary of GSEA evaluating Hallmark gene sets within the MSigDB v7.0 database upon ZBED2 cDNA expression in 15 PDA cell lines. NES scores for each gene set were averaged across each cell line and this value was used to rank each gene set. The top three most repressed (top panel) and activated (bottom panel) gene sets are shown. “#” indicates the number of cell lines in which the indicated gene set was determined to be repressed or activated with a false discovery rate (FDR) or <0.1. NES, normalized enrichment score; SD, standard deviation. (*B*) RT-qPCR analysis of the ISGs *STAT2, MX1* and *CMPK2* in HPAFII cells (left panel) and PANC0403 cells (right panel) following infection with ZBED2 cDNA or the empty vector. (*C* and *D*) Western blot analysis of HSC70 and ZBED2 (*C*) and RT-qPCR analysis of *ZBED2*, *STAT2* and *CMPK2* (*D*) in PANC0403 cells following infection with ZBED2 shRNAs or a control shRNA (shNEG). (*E* and *F*) RT-qPCR analysis of *IRF1*, *MX1* and *ZBED2* in PANC0403 cells following CRISPR-based targeting of IFNAR1 with three independent sgRNAs or a control sgRNA (*E*) and subsequent treatment with 20ng/µL of IFNβ for 12 hours (*F*). (*G*) *ZBED2* expression across all cancer cell lines within the CCLE database (left panel) or in the indicated tissue lineages stratified according to high or low *ZBED2* expression. Each cell line is depicted as a single dot. (*H*) RT-qPCR analysis of *IRF1* and *ZBED2* following treatment with 20ng/µL of IFN-β or IFN-γ for 12 hours in PDA cell lines. Mean+SEM is shown. n=3. ns, not significant, \**p*<0.05, **p <0.01, ***p <0.001 by Student’s t test.

**Fig. S3.**
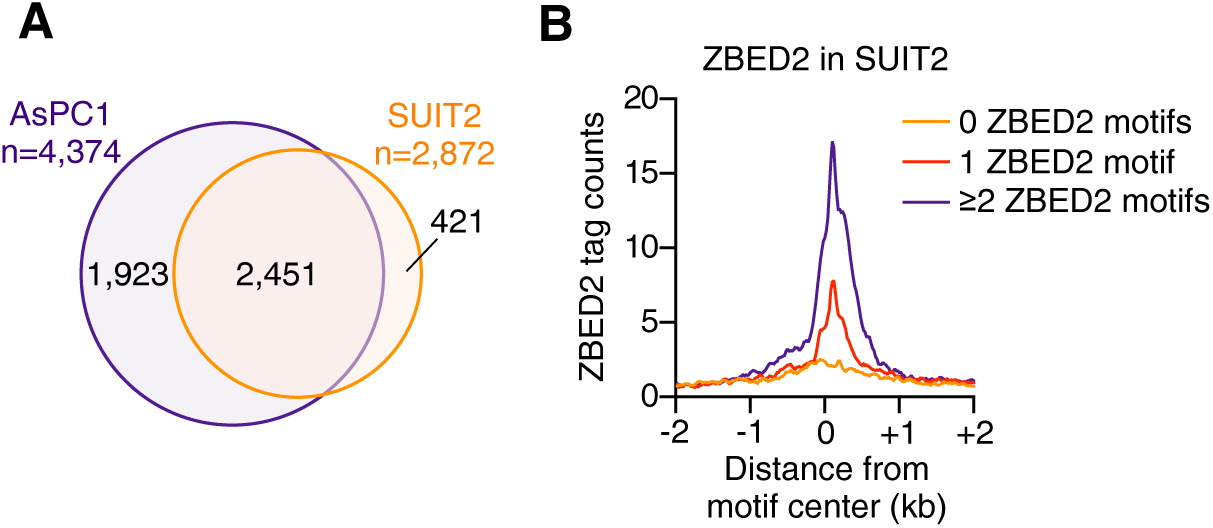
Epigenomic analysis of ZBED2. Related to Fig. 3. (*A*) Overlap of FLAG-ZBED2 ChIP-seq peaks in AsPC1 and SUIT2 cells. (*B*) Meta-profile comparing ZBED2 occupancy in SUIT2 cells around the center of peaks with 0, 1 or ≥2 motif counts. ZBED2 binding intensity is shown as sequencing depth normalized tag count.

**Fig. S4.**
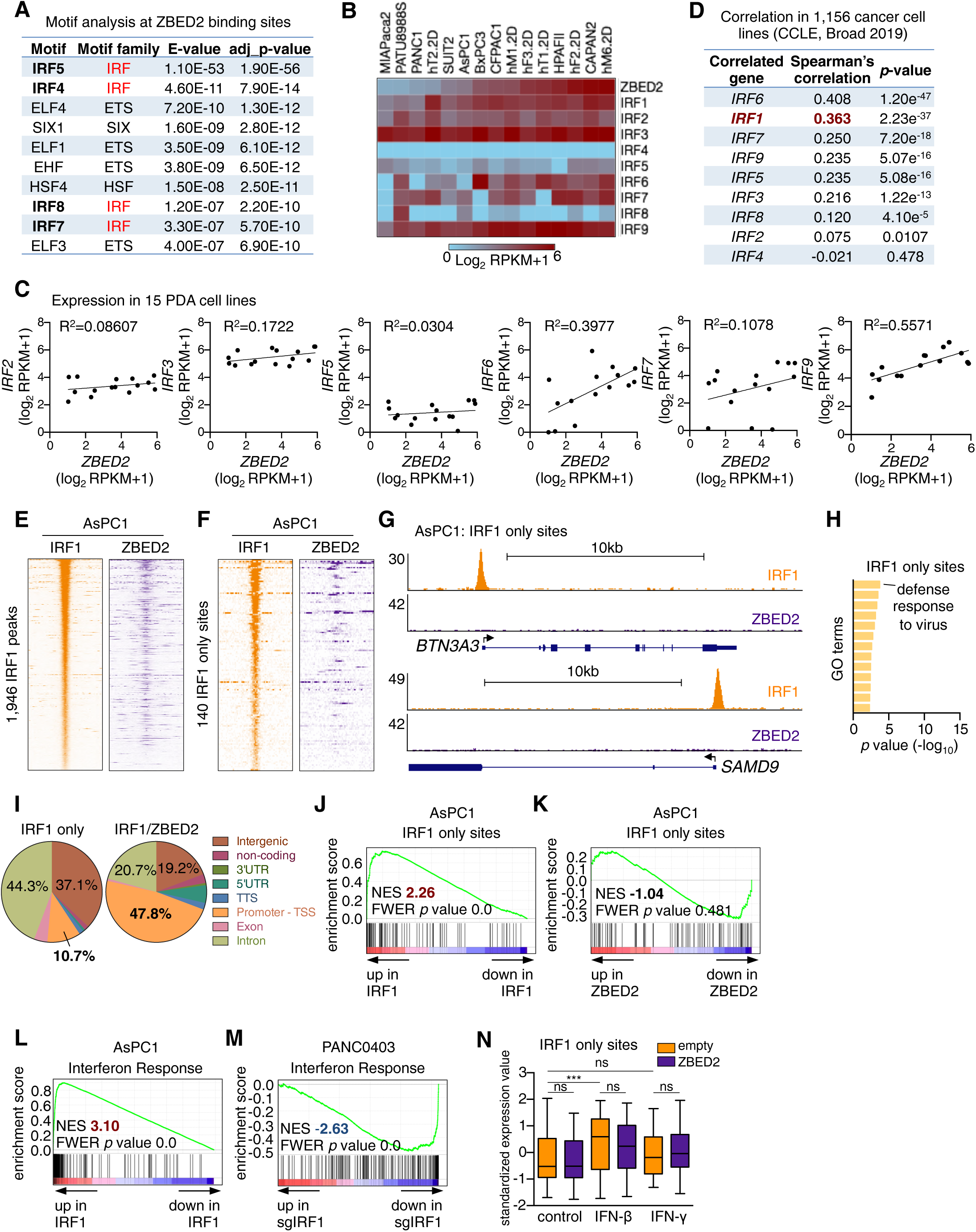
Antagonistic regulation of ISG promoters by ZBED2 and IRF1. Related to Fig. 4. (*A*) Summary of CentriMo motif enrichment analysis for JASPAR motifs at ZBED2 binding sites. The top 10 transcription factor (TF) motifs ranked by E value are shown. The nucleotide sequence 200bp up- or downstream of the peak summit of the top 1000 ZBED2 peaks in AsPC1 cells was used for this analysis. (*B*) Expression of *ZBED2* and IRF family TF genes in 15 human PDA cell lines. (*C*) *ZBED2* expression versus *IRF2*, *IRF3*, *IRF5*, *IRF6*, *IRF7* and *IRF9* expression across 15 human PDA cell lines. (*D*) *ZBED2* expression correlation with IRF family TF genes in 1,156 cancer cell lines from the CCLE database analyzed using CBioPortal. (*E* and *F*) Density plots showing IRF1 and FLAG-ZBED2 enrichment surrounding a 2-kb interval centered on the summit of all IRF1 peaks (*E*) or 140 random IRF1 peaks that do not intersect with FLAG-ZBED2 sites in AsPC1 cells (*F*), ranked by IRF1 peak intensity. (*G*) ChIP-seq profiles of IRF1 and FLAG-ZBED2 in AsPC1 cells at the promoters of *BTN3A3* and *SAMD9*. (*H*) Gene ontology (GO) analysis with Metascape of genes annotated by HOMER to IRF1 only sites. Terms are ranked by their significance (*p* value) and no terms reached the significant threshold (-log_10_ *p* value >12). (*I*) Pie chart showing the distribution of 140 IRF1 only peaks (left) or IRF1/ZBED2 peaks (right) in AsPC1 cells. TTS, transcription termination site; TSS, transcription start site; UTR, untranslated region. (*J* and *K*) GSEA plots evaluating protein coding genes annotated by HOMER to IRF1 only sites upon IRF1 (*J*) or ZBED2 (*K*) cDNA expression in AsPC1 cells. (*L* and *M*) GSEA plots evaluating the Interferon Response signature upon IRF1 cDNA expression in AsPC1 cells (*L*) or IRF1 knockout in PANC0403 cells (*M*). (*N*) Expression levels of protein coding genes annotated to IRF1 only sites following 12-hour treatment with 0.2ng/µl of IFN-β, IFN-γ or control. ***p<0.001, ns = not significant, by one-way ANOVA.

**Fig. S5.**
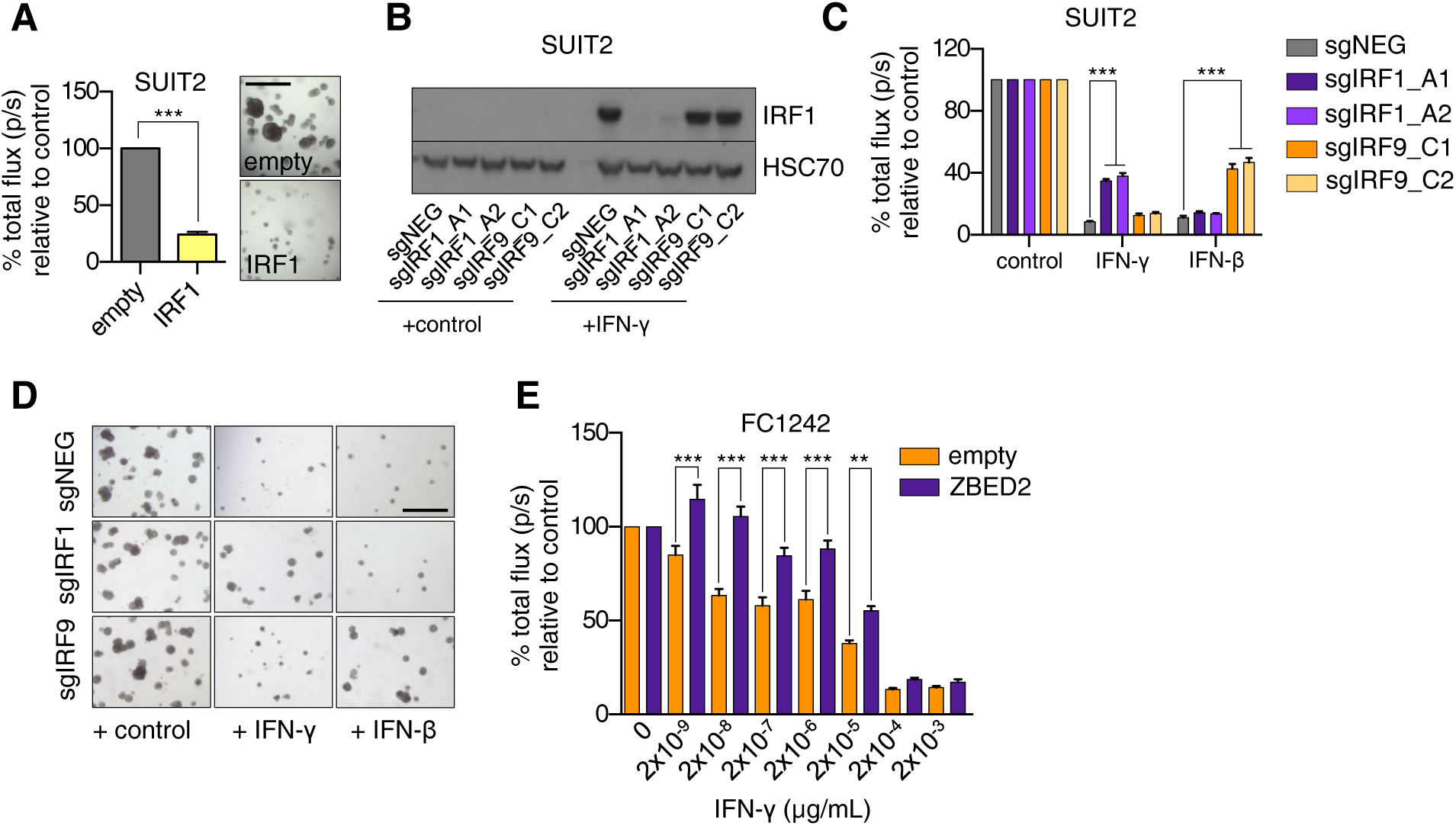
ZBED2 protects PDA cells from IRF1- and interferon-γ-induced growth arrest. Related to Fig. 5. (*A*) Luciferase-based quantification of cell viability of SUIT2 cells grown in Matrigel on day 7 post infection with IRF1 cDNA or the empty vector. Representative bright field images (right panel) are shown. Scale bar indicates 200µm. (*B-D*) SUIT2 cells infected with sgRNAs targeting IRF1, IRF9 or a control sgRNA (sgNEG) were plated in Matrigel and grown for 7 days in the presence of 20ng/µl of IFN-γ, IFN-β or control. Representative western blots following overnight stimulation with 20ng/µl of IFN-γ (*B*), luciferase-based quantification (*C*) and representative bright field images on day 7 of the assay (*D*) are shown. Scale bar indicates 500µm. (*E*) FC1242 KPC cells were infected with the ZBED2 cDNA or an empty vector and grown in Matrigel with the increasing concentrations of IFN-γ. Bar chart shows luciferase-based quantification on day 7. Mean+SEM is shown. n=3. For (*A*) and (*C*), ***p <0.001 by Student’s t test. For (*E*), ***p<0.001, **p<0.01 by one-way ANOVA.

**Fig. S6.**
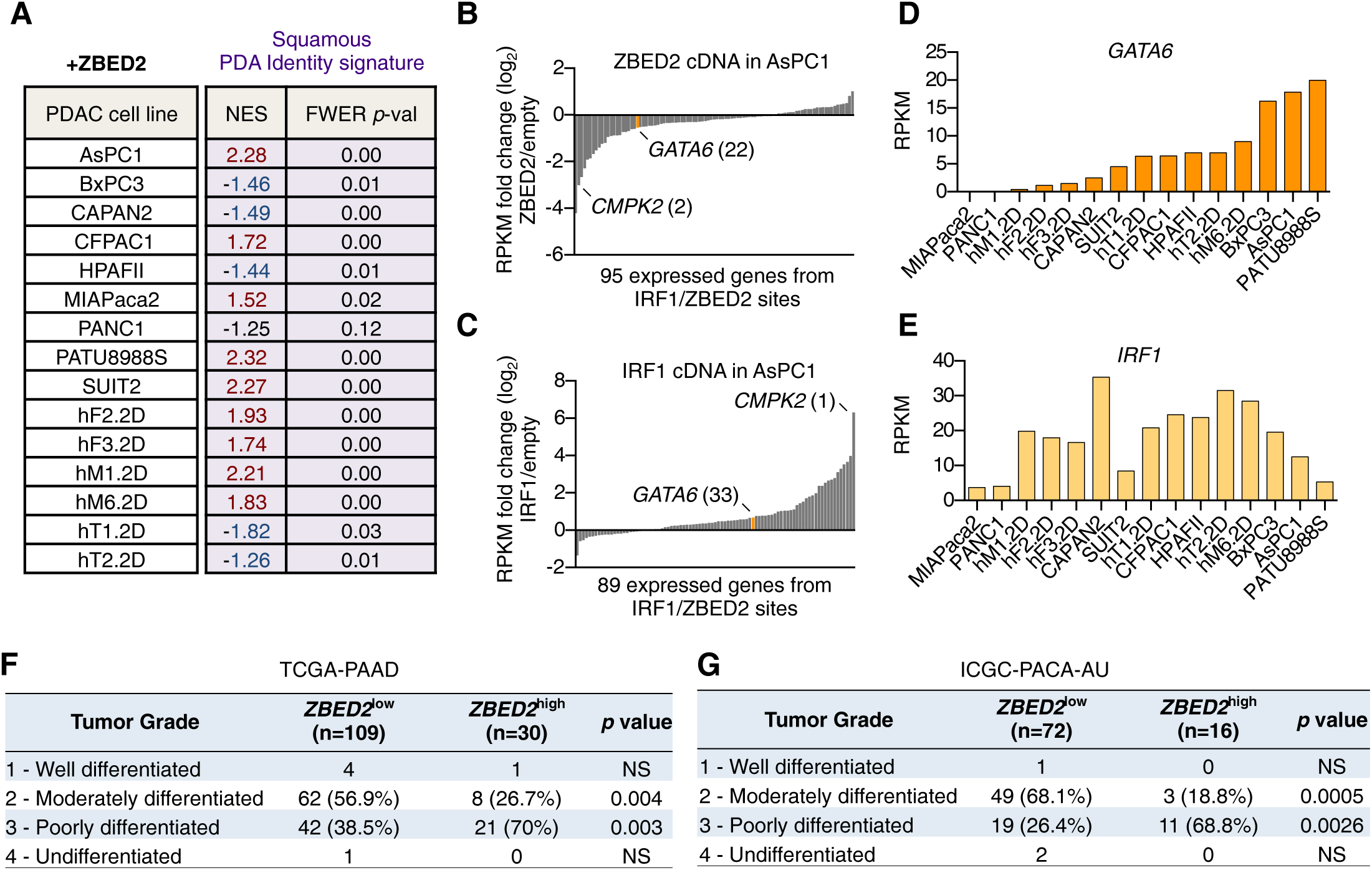
ZBED2 represses pancreatic progenitor lineage identity in PDA. Related to Fig. 6. (*A*) Summary of GSEA evaluating the Squamous PDA Identity signature upon ZBED2 cDNA expression in 15 PDA cell lines. (*B* and *C*) Expression changes at IRF1/ZBED2 bound genes in AsPC1 cells infected with ZBED2 cDNA (*B*) or IRF1 cDNA (*C*) versus those infected with an empty vector control. *GATA6* and *CMPK2* are labeled along with their rank with respect to downregulated (*B*) or upregulated (*C*) genes. (*D* and *E*) *GATA6* (*D*) and *IRF1* (*E*) expression in 15 human PDA cell lines. (*F* and *G*) Proportion of PDA patient samples from the indicated studies stratified as *ZBED2*^low^ or *ZBED2*^high^ classified based on their tumor differentiation status (grade). Statistical significance for the indicated comparisons was assessed using Fisher’s Exact Test, ns = not significant.

## Supplementary Datasets

Overview of supplementary datasets

Dataset. S1. Ranked list of expressed transcription factors in primary PDA tumor samples versus normal pancreas tissue.

Dataset. S2. *ZBED2* expression values as normalized counts in normal and neoplastic pancreatic organoids.

Dataset. S3. Results from GSEA for gene sets within the MSigDBv7.0 following ZBED2 knock out in PANC0403 or ZBED2 cDNA expression in HPAFII cells.

Dataset. S4. Interferon Response signature genes.

Dataset. S5. Results from GSEA for Hallmark gene sets following ZBED2 cDNA expression in 15 PDA cell lines.

Dataset. S6. 2,451 high confidence ZBED2 peaks and their HOMER annotations.

Dataset. S7. GSEA of ZBED2 bound genes following ZBED2 cDNA expression in HPAFII cells. Dataset. S8. Results from ZBED2 motif scanning at human promoters.

Dataset. S9. 1,946 high confidence IRF1 peaks and their HOMER annotations.

Dataset. S10. 140 IRF1/ZBED2 sites and the control set of IRF1 only sites with their HOMER annotations.

Dataset. S11. Fold change in IRF1 tag counts in AsPC1-ZBED2 versus AsPC1-empty cells at high confidence IRF1 sites with their HOMER annotations.

Dataset. S12. Ranked lists of expressed TFs in Basal-like versus Classical or Squamous versus Progenitor PDA tumor samples from three independent studies.

Dataset. S13. Sequences for RT-qPCR primers, sgRNAs and shRNAs used in this study.

Additional information is provided as a header in the first page of each Excel workbook.

